# WNT7B drives a program for pancreatic cancer subtype switching and progression

**DOI:** 10.1101/2024.12.18.628910

**Authors:** J. Sprangers, J.M. Bugter, D. Xanthakis, M. Boekhout, R.N.U. Kok, L.A.A. Brosens, F. Dijk, M.J. Rodríguez Colman, J.Y. van der Vaart, M.M. Maurice

**Affiliations:** Oncode Institute, The Netherlands; Center for Molecular Medicine, UMC Utrecht, Utrecht, The Netherlands; Department of Pathology, University Medical Center Utrecht, Utrecht University, 3584 CX Utrecht, the Netherlands; Department of Gynaecologic Oncology, Amsterdam University Medical Centres, Amsterdam, The Netherlands; Department of Pathology, Amsterdam University Medical Centres, Amsterdam, The Netherlands

## Abstract

Hyperactivation of WNT signaling is a well-established hallmark of cancer. Various epithelial cancers express high levels of WNT7B and WNT10A that are not commonly expressed during tissue homeostasis, but rather associate with tissue development and regeneration. Although increased WNT7B/10A expression correlates with aggressive disease and lower patient survival rates, the mechanism by which these WNTs influence cancer progression remains unknown. Here, we use patient-derived organoids to show that tumor-intrinsic expression of WNT7B/10A drives survival and growth of advanced pancreatic ductal adenocarcinoma (PDAC). Bulk and single-cell profiling reveal that WNT7B drives proliferation and promotes expression of a poor prognosis basal-like state by preventing expression of a more differentiated, classical PDAC signature. By generating WNT7B reporter organoids, we show that heterogeneously distributed WNT-high PDAC cells are shifted towards a more basal-like phenotype and stably co-exist with WNT-low/negative lineages. Furthermore, hybrid co-cultures of WNT7B-proficient and -knockout PDAC organoids demonstrate that WNT-sending cells drive survival and proliferation of neighboring WNT-negative cells within the cancer epithelium via short range, cell contact-dependent signaling. In summary, our work uncovers a prominent role of WNT7B/10A in driving PDAC subtype heterogeneity and argues that WNT inhibition may be applied to force a ‘class switch’ to a more differentiated, less aggressive cancer subtype that correlates with improved therapeutical response.

## Introduction

Pancreatic ductal carcinoma (PDAC) is one of the most lethal malignancies, characterized by treatment-resistance and high rates of metastasis (Dyba *et al*., 2021). Mortality rates are further increased due to late detection and limited overall treatment options. PDAC comprises the most common form of pancreatic cancer that originates from the exocrine compartment of the pancreas. PDAC formation may arise from various preneoplastic stages, including pancreatic intraepithelial neoplasms (PanINs) or more mucinous-type lesions such as intraductal papillary mucinous neoplasms (IPMNs) and mucinous cystic neoplasms (MCNs) (Distler *et al*., 2014). Commonly identified driver mutations found in PDAC tissues comprise mutations in components of the EGF/RAS, P53 and WNT pathways (Bailey *et al*., 2016). Over the last decade, various studies aimed to classify PDAC tumors into subtypes based on the expression of transcriptional programs (Bailey *et al*., 2016; Chan-Seng-Yue *et al*., 2020; Moffitt *et al*., 2015). Collectively, two consensus subtypes are recognized and referred to as classical and basal-like PDAC. PDAC tumors of the classical type are enriched for transcriptional programs that are linked to differentiation and development, and these tumors generally display a better response to standard-of-care chemotherapy (Aung *et al*., 2018; Robertson *et al*., 2024; Suurmeijer *et al*., 2022). By contrast, basal-like tumors represent a more aggressive subtype with lower overall survival rates characterized by activation of gene networks in inflammation, metabolic reprogramming and hypoxia (Bailey *et al*., 2016; Brunton *et al*., 2020). Despite an estimated prevalence of around 20% for basal-like tumors (Zhou *et al*., 2021; Suurmeijer *et al*., 2022; Williams *et al*., 2023), recent findings indicate extensive subtype heterogeneity within PDAC tumors, which challenges straightforward binary subtype classification (Krieger *et al*., 2021; Raghavan *et al*., 2021; Saillard *et al*., 2023; Williams *et al*., 2023).

Although downstream WNT pathway mutations, exemplified by mutations in *APC* or *β-catenin*, are generally lacking in PDAC (Bugter *et al*., 2021), a substantial subset of tumors displays features of elevated WNT signaling activity. This is mostly attributed to mutations in the WNT pathway negative feedback regulator *Ring finger protein 43* (*RNF43)*, at reported frequencies between 5-10% in PDAC (Bailey *et al*., 2016; Collisson *et al*., 2019; Waddell *et al*., 2015). *RNF43* encodes for a membrane-bound E3 ubiquitin ligase that targets Frizzled (FZD) proteins, the primary receptors for WNT, for endocytosis and lysosomal degradation (Hao *et al*., 2012; Koo *et al*., 2012). Mutational inactivation of *RNF43* leads to an overabundance of WNT receptors at the cell surface, which confers a WNT hypersensitive growth state to cancer cells (Bugter *et al*., 2021; Koo *et al*., 2012). In another strategy to identify WNT-addicted pancreatic cancers, cultured PDAC organoids that grow without an exogenous WNT source displayed strong growth inhibition when treated with Porcupine inhibitors (Seino *et al*., 2018), small molecules that potently inhibit WNT secretion (Liu *et al*., 2013). Together, these findings indicate that a substantial fraction of PDAC tumors displays a growth state that relies on WNT ligand production by the cancer epithelium itself.

In line with the above, a subset of PDAC tumors displays elevated expression of various WNTs, with WNT7B and WNT10A as most prominent examples (Arensman *et al*., 2014; Brunton *et al*., 2020; Seino *et al*., 2018). Strikingly, WNT7B plays a key role during pancreatic development (Afelik *et al*., 2015), while WNT signaling is dispensable for adult pancreas maintenance (Keefe *et al*., 2012; Murtaugh *et al*., 2005). In addition, re-induced epithelial expression of both Wnt7b and Wnt10a in mice was observed upon damage of various epithelial tissues, including lung (Nabhan *et al*., 2018; Volckaert *et al*., 2017), liver (Okabe *et al*., 2016; Planas-Paz *et al*., 2019) and intestine (Berger *et al*., 2016; Tan *et al*., 2021), indicating that these WNTs regulate a conserved program for epithelial tissue repair. Besides their pronounced expression in PDAC, WNT7B and WNT10A also display elevated expression in other malignancies, including prostate cancer (Zheng *et al*., 2013), bile duct cancer (Boulter *et al*., 2015) and esophageal cancer (Werner *et al*., 2023). Notably, WNT7B and WNT10A expression levels generally correlate with more aggressive subtypes and, consequently, a lower survival probability of patients (Brunton *et al*., 2020; Hsu *et al*., 2012; Liu *et al*., 2022; Long *et al*., 2015; Seino *et al*., 2018). This correlation was validated for pancreatic cancer, where increased expression of WNT7B and WNT10A links to the more aggressive basal-like subtype (Arensman *et al*., 2014; Brunton *et al*., 2020; Raghavan *et al*., 2021; Seino *et al*., 2018). Despite this clear link of high WNT7B and WNT10A expression with poor prognosis cancer subtypes, an in-depth understanding of how these WNTs drive cancer development and progression is lacking.

In this study, we employ human patient-derived PDAC organoids to examine how WNT7B- and WNT10A-activated programs drive PDAC growth and progression. Using genetic editing strategies and single cell transcriptional approaches, we uncover that these WNTs are expressed by a subset of epithelial tumor cells to mediate short range, cell contact-dependent signaling that drives survival and growth of all tumor cells. Furthermore, our results reveal that WNT7B/10A-mediated signaling suppresses a differentiated, classical gene expression profile, and thus cause a relative increase in the basal-like expression profile that is most prominent within the WNT-high fraction of PDAC tumor cells. Together, our data reveal that WNT7B is a functional marker of the poor prognosis-linked basal-like transcriptional state.

## Results

### A subset of basal-like PDAC tumors expresses high levels of WNT7B and WNT10A

To investigate WNT expression levels in PDAC, we examined transcriptome data of tumor tissues derived from a large cohort of pancreatic cancer patients (Suurmeijer *et al*., 2022). PurIST scoring was used to assess tumor subtype state (Rashid *et al*., 2020). WNT2, WNT5A and WNT11 were prominently expressed across all PDAC patient tumor tissue subsets (Figure S1A). By contrast, expression of WNT7A, WNT7B and WNT10A, as well as Porcupine (PORCN), an essential acyltransferase for WNT secretion, were found enriched within basal-like annotated tumors (Figure 1A,B, Figure S1A,B). Of note, the cellular origin of WNTs (tumor, stroma or associated cell types) could not be determined in this dataset.

**Figure 1.**
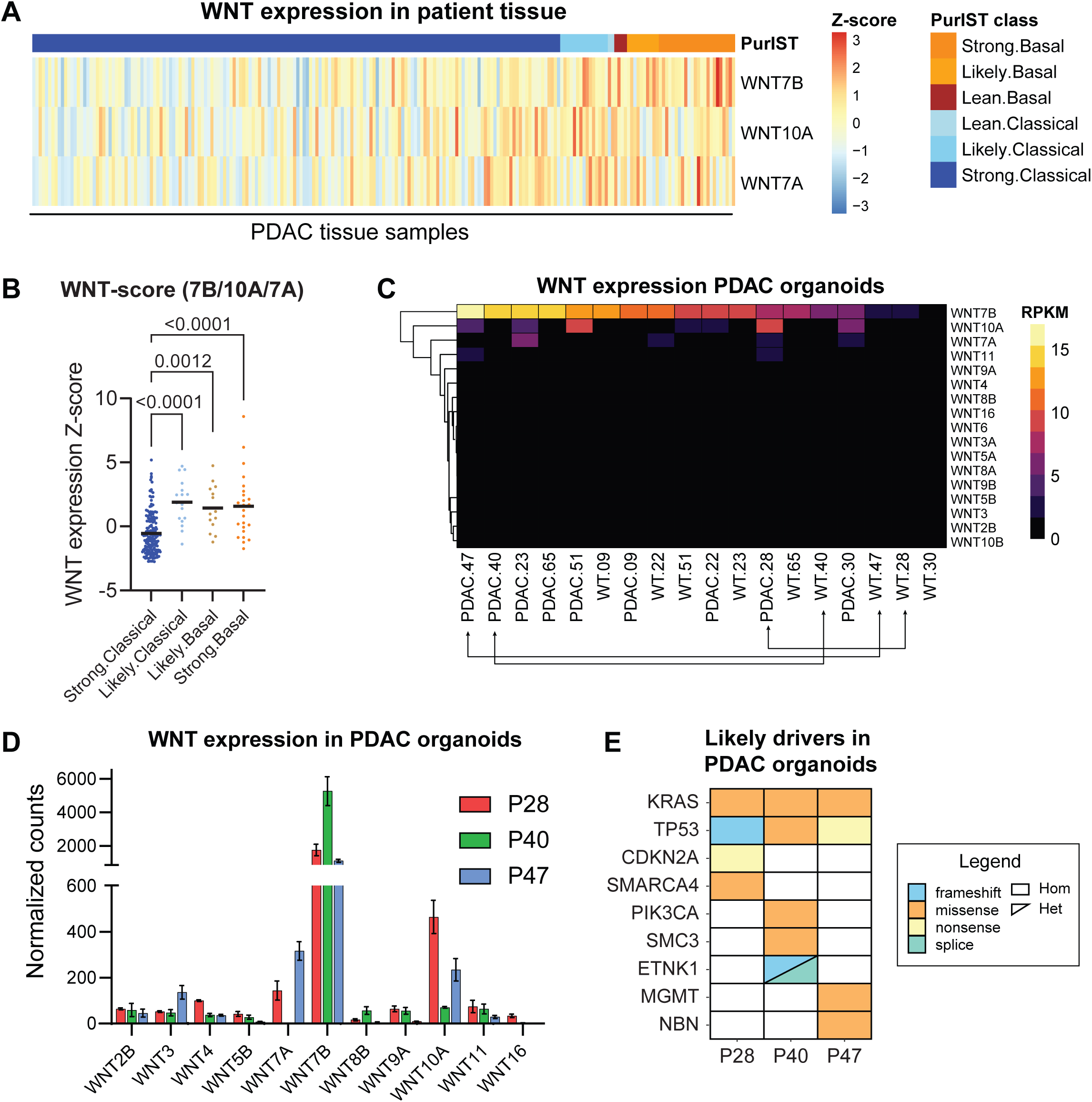
Subset of PDAC organoids displays increased WNT protein expression. (A) Heatmap of bulk tumor tissue samples within the SPACIOUS-II cohort. PurIST subtype scoring is indicated for each sample in the top row. (B) Dot plot showing PurIST score of each PDAC sample, categorized by inferred PurIST subtype, versus a cumulative score of epithelial WNT ligands (WNT7A/7B/10A). P-values of Kruskal-Wallis test indicated. (C) Heatmap showing RNA expression (RPKM) of epithelial WNTs in organoids derived from PDAC patients. (D) Bulk RNA sequencing derived WNT expression in selected PDAC organoid lines, displayed as DEseq2 normalized counts. (E) Oncoprint plot showing likely genetic driver mutations in each PDAC organoid line. Hom: homozygous mutation; Het: heterozygous mutation.

To examine a potential driver role of epithelial PDAC-associated WNTs, we screened a library of patient-derived PDAC organoid lines for WNT expression (Driehuis *et al*., 2019). Strikingly, WNT7B, WNT10A and WNT7A comprised the most abundantly expressed WNTs in PDAC lines in comparison to healthy pancreatic organoids (Figure 1C). Three patient-derived PDAC organoid lines with high WNT7B and/or WNT10A expression (P28, P40 and P47) were selected for further investigation (Figure 1C, Figure S1C). While P40 and P47 displayed a cystic appearance, P28 organoids exhibited a more dense, spherical morphology, similar to earlier reports (Krieger *et al*., 2021; Seino *et al*., 2018) (Figure S1D). WNT7B was validated as the most prominent WNT expressed in all three PDAC organoid lines, followed by lower levels of WNT10A and WNT7A in P28 and P47 (Figure 1D, Figure S1E). Since organoids are epithelial-only cultures, these data indicate that WNT7A/7B/10A are generated by cancer epithelial cells, while other prominent PDAC-associated WNTs (WNT2, WNT5A and WNT11) are produced by extrinsic, cancer-associated cell types. DNA sequencing of all three organoid lines revealed the presence of common PDAC-associated genetic driver mutations, including mutant *KRAS* and *TP53*, while mutations in known WNT pathway components, such as *APC*, *CTNNB1*, *AXIN1* and *RNF43* (Bugter *et al*., 2021), were absent (Figure 1E). Thus, this subset of PDAC organoids acquired the capacity to display increased WNT7B/10A expression in the absence of other mutations that promote WNT pathway activation or hypersensitivity.

### Cancer epithelium-intrinsic WNTs are essential for PDAC organoid growth and survival

Next, we examined whether PDAC organoid growth and survival depends on tumor cell-derived WNTs, by treating organoids with Porcupine inhibitor LGK-974 (LGK) that potently blocks secretion of WNT proteins (Liu *et al*., 2013). Of note, organoid lines were established and cultured in medium that lacks WNT but includes R-spondin. R-spondin binds to LGR5 and RNF43, blocking RNF43 function and subsequently increasing FZD-mediated WNT signaling (Hao *et al*., 2012; Zebisch *et al*., 2013). All three organoid lines were highly sensitive to 7 days of LGK treatment as shown by their failure to grow out after the first passage (Figure 2A, B). This phenotype could be rescued in all lines by supplementation with WNT3A-conditioned medium (WNT3A-CM), but not by L-cell conditioned control medium (LCM, Figure 2B, Figure S2A). In addition, WNT pathway activation mediated by WNT surrogate (WNTsur), a bispecific protein that heterodimerizes Frizzleds and the WNT co-receptor LRP5/6 (Janda *et al*., 2017), also rescued LGK-induced growth arrest (Figure 2B, Figure S2A). Thus, the growth and survival of selected PDAC organoids are reliant on WNT pathway activation induced by self-produced WNTs.

**Figure 2.**
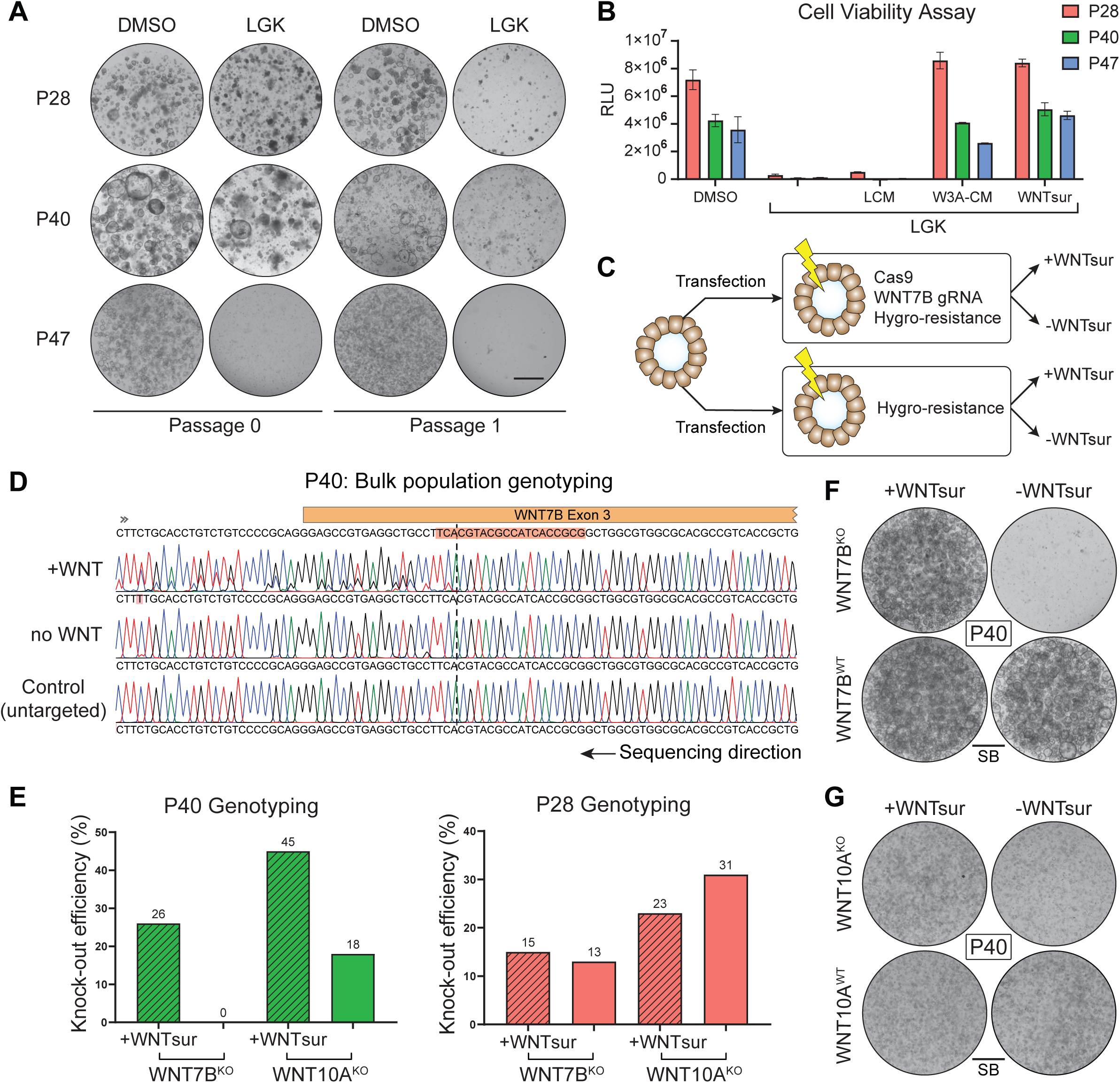
PDAC organoids depend on endogenous WNT proteins. (A) Brightfield images of DMSO and LGK-treated PDAC organoids at indicated concentrations before and after passaging. (B) Cell viability assay of DMSO and LGK-treated PDAC organoids with indicated rescue conditions (passage 1, day 7). (C) Schematic of electroporation procedure for WNT knock-out generation in PDAC organoids. (D) Genotyping of targeted WNT7B locus in bulk P40 organoid populations, with rescue conditions indicated. Guide sequence for WNT7B targeting highlighted in orange, cut site indicated with dashed line. (E) WNT7B knock-out efficiency scores of targeted bulk P28 and P40 organoid populations. (F-G) Brightfield images of outgrowth experiments with P40 WNT7B (F) and WNT10A (G) knock-out clonal lines in presence or absence of exogenous WNT at passage 1 day 7 (WNT7B clones) and passage 1 day 5 (WNT10A clones). LCM: L-cell conditioned medium; W3A-CM: WNT3A-conditioned medium. RLU: relative luciferase units. Scale bar (SB) = 1000µm.

To investigate how specific PDAC-derived WNTs contribute to organoid growth and survival, we generated WNT-specific knock-out lines using P40 organoids that mainly produce WNT7B (Figure 1E). After introducing Cas9 and WNT7B-targeting gRNA constructs, we split the targeted cells and supplemented one population with WNTsur, while leaving the other population without WNT, similar to non-targeted cell lines (Figure 2C). Subsequent genotyping revealed that knock-out traces within the WNT7B locus were only detected in WNTsur-supplemented conditions (knock-out score 26%), indicating that the outgrowth of WNT7B-knockout organoids depends on an exogenous WNT source (Figure 2D, E). Additionally, these data reveal that WNT7B knock-out cells are not rescued by untargeted neighboring WNT7B-secreting organoids, suggesting that WNT7B does not diffuse in extracellular space (Figure S2B). Of note, attempts to knockout WNT10A in P40 organoids did not affect outgrowth of these organoids, likely because remaining levels of WNT7B are sufficient to rescue loss of WNT10A in this organoid line (Figure 2E).

To assess whether WNT7B loss renders P40 organoids fully dependent on external WNT, we generated clonal lines carrying homozygous loss-of-function mutations in WNT7B (WNT7B^KO^) or WNT10A (WNT10A^KO^) and grew them out in the presence of WNTsur. Genotyping confirmed that premature stop codons were introduced in both alleles (Figure S2C-F). Notably, removal of WNTsur from the culture medium of WNT7B^KO^ organoids led to a complete growth arrest, while WNT7B^WT^ organoids that went through the same procedure, remained unaffected by WNTsur removal (Figure 2F). By contrast, WNT10A^KO^ knockout organoids still grew out upon removal of exogenous WNT (Figure 2G). Collectively, these data show that the growth and survival of P40 PDAC organoids fully depends on the production of WNT7B by cancer epithelial cells.

Next, we investigated the dependency of P28 PDAC organoids on the production of individual WNTs. P28 PDAC organoids generate a broader set of WNTs as compared to P40, with highest expression of WNT7B, WNT10A and WNT7A (Figure 1E, S1D). Using the same procedure as for P40 (Figure 2C), we generated single and double knock-out P28 clones for WNT7B and WNT10A. All P28-derived WNT knock-out clones, including a WNT7B/10A double knock-out clone, were able to grow out in absence of supplementation with exogenous WNT (Figure 2E, S2G-H). At the same time, all P28 clonal variants remained LGK-sensitive (Figure S2H), indicating that the low expression levels of other epithelial WNTs, such as WNT7A, can rescue growth and survival of these PDAC cells. Thus, a WNT-mediated growth and survival program in the PDAC epithelium can be mediated by only one, or multiple WNTs.

### PDAC-derived WNTs promote a basal-like PDAC subtype cellular state

To assess the immediate downstream effects of epithelial WNTs that mediate PDAC organoid survival, we treated all three organoid lines for 24h with LGK and examined alterations in transcriptional profiles using RNA sequencing (Figure 3A). At this early timepoint after start of LGK treatment, alterations in organoid morphology and growth were not yet visible (Figure S3A). To identify shared principles of WNT-mediated signaling within the PDAC epithelium, we focused on differentially expressed genes (DEGs) in two or more organoid lines (Figure 3B, Table S1). GO term analysis showed that downregulated gene sets were linked to WNT/β-catenin signaling and DNA replication, thus confirming efficient WNT inhibition at 24h after treatment (Figure 3C). In agreement, core WNT/β-catenin target genes were downregulated, including AXIN2, LGR5, RNF43 and ZNRF3 (Figure 3D-E, S3B) (Boonekamp *et al*., 2021). Furthermore, several gene sets associated with previously described WNT10A- and WNT7B-driven developmental processes, including tooth and bone development, were significantly downregulated in LGK-treated PDAC organoids (Figure 3C) (Song *et al*., 2020; Wang *et al*., 2022; Xu *et al*., 2017). These data thus reveal that PDAC-derived WNTs drive canonical WNT signaling and cell proliferation, as well as conserved gene programs linked to tissue development.

**Figure 3.**
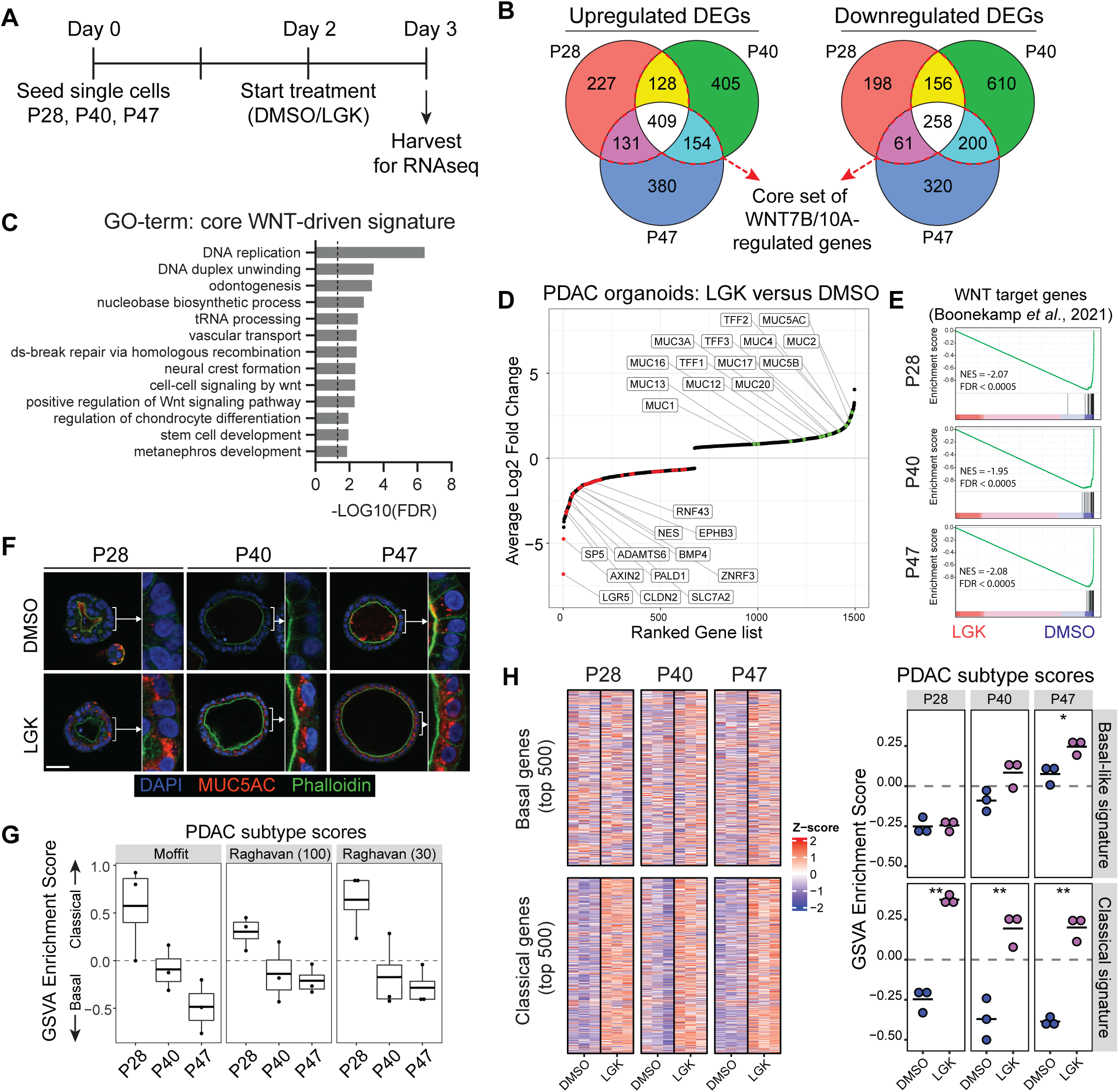
WNT proteins drive PDAC cell proliferation and suppress classical subtype gene expression. (A) Schematic overview of experimental setup for bulk RNA sequencing. (B) Overlapping up- and downregulated differentially expressed genes (DEGs) between PDAC organoid lines (FC>1.5; FDR<0.05). (C) Biological process GO-term analysis on all significantly downregulated DEGs after LGK-974 treatment (DEGs were included when average expression over all samples >50). Selected GO-terms of different hierarchies are displayed. Dashed line indicates significance threshold of FDR=0.05. (D) Dot plot of significantly DEGs, ranked by increasing foldchange. Annotated genes are WNT target genes (red) or *Mucin*/*TFF* genes (green). (E) GSEA analysis of core WNT target genes (Boonekamp et al., 2021). (F) MUC5AC staining in PDAC organoids after 48h of DMSO- or LGK-treatment. Zoom-ins of indicated organoid images is displayed. (G) GSVA analysis of PDAC subtype signatures of indicated gene set size (Moffitt et al., 2015; Raghavan et al., 2021). (H) Heatmap (left) and GSVA analysis (right) of top 500 basal-like and classical gene signatures in DMSO- and LGK974-treated conditions. Asterisks indicate p-values (t-test, P28: p=0.0021; P40: p=0.0047; P47 classical: p=0.0029; P47 basal-like: p=0.02). SB=50µm; Ds: double strand.

Conversely, LGK treatment induced upregulation of several genes linked to pathways of cell differentiation, including the production of multiple mucins (MUC5AC, MUC2) and Trefoil Factors (TFF1, TFF2, TFF3) (Figure 3D). We confirmed these findings at the protein level, by showing increased staining of MUC5AC in LGK-treated organoids (Figure 3F). Together, these genes encompass known signatures for gastric metaplasia (MUC5AC, TFF1, ANXA10) and intestinal metaplasia (MUC2, REG4), that are commonly activated in context of pancreatic injury as well as in preneoplastic lesions of PDAC, including PanIN and IPNM (Guppy *et al*., 2012; Schlesinger *et al*., 2020; Strobel *et al*., 2010; Tsai *et al*., 2015; Zhang *et al*., 2021). Notably, several upregulated DEGs comprise secretory genes that are not expressed in healthy pancreatic tissue, including MUC5AC and MUC2 (Wang *et al*., 2020). In line with these findings, gene sets associated with gastric metaplasia and secretory cell types, including gastric pit cells and goblet cells, are enriched in LGK-treated organoids (Figure S3D-E, Table S2) (Busslinger *et al*., 2021; Chen *et al*., 2021).

Since WNT expression is correlated with the basal-like PDAC subtype state (Brunton *et al*., 2020; Zhong *et al*., 2022), we used our transcriptional dataset to infer PDAC organoid subtype state and examine whether WNT-induced programs play a role in determining these signatures. When comparing subtype scores using previously published PDAC subtype signatures (Moffitt *et al*., 2015; Raghavan *et al*., 2021) we observe a higher classical score for P28 and P40, while P47 is more basal-like (Figure 3G). Next, we assessed how LGK-mediated WNT inhibition alters the expression of these PDAC subtype signatures. Classical subtype-linked genes were increased in all LGK-treated organoid lines, while genes associated with the basal-like PDAC subtype remained largely unaffected (Figure 3H). Taken together, our data argue that PDAC-derived WNTs suppress the transcriptional program of the classical PDAC subtype while maintaining basal-like gene expression, leading to a relative shift towards the basal-like subtype.

### PDAC-derived WNTs are heterogeneously expressed within PDAC organoids

To study how epithelial WNT expression is organized at the single cell level within PDAC organoids, we performed single cell RNA sequencing analysis. As expected, cellular profiles from all three organoid lines cluster separately, due to differences between patients (Figure S4A) (Krieger *et al*., 2021). To allow for the identification of shared principles, we performed data integration using the Seurat package (see methods), which reveals multiple cell clusters that share characteristics within all three PDAC organoid lines (Figure 4A). Gene expression analysis indicated that these clusters correspond with cycling cells (Cycling-S, Cycling-G2/M), more differentiated-like cancer cells (Diff1, Diff2), as well as cell types enriched for genes involved in cilia formation (Ciliated) or genes linked to an activated interferon response (IFN cells) (Figure 4A, S4B-D). The two differentiated-like cell populations that reside in G1 show partial overlap with gene expression from cycling clusters (Diff1) (Figure S4E), or display expression of genes linked to cell adhesion, hypoxia and cholesterol biosynthesis (Diff2) (Figure S5A). One cluster of cells that carried a low number of reads and genes per cell, termed low quality (‘LowQ’), was disregarded for further analysis (Figure 4A, S4B).

**Figure 4.**
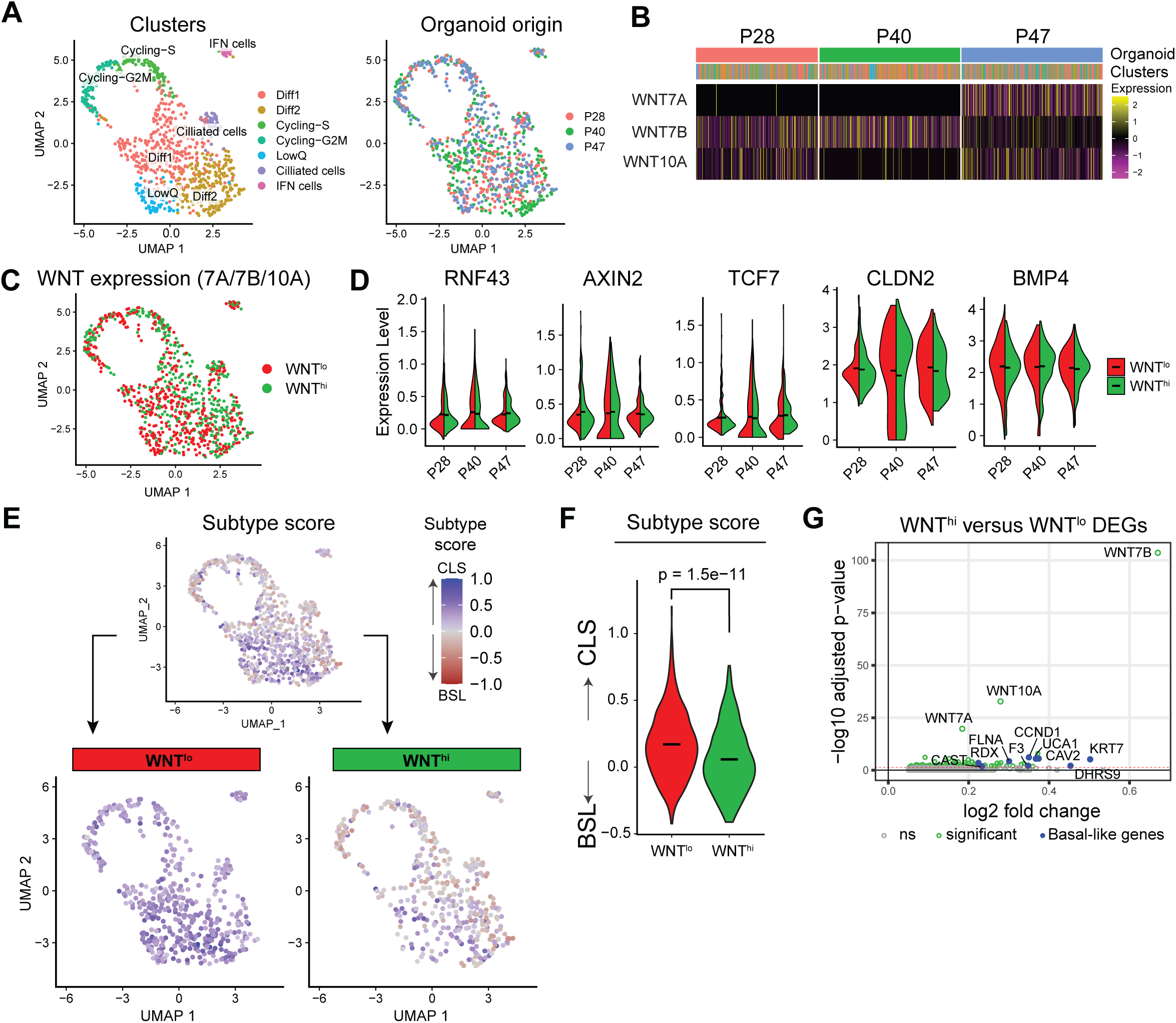
Epithelium-derived WNTs are heterogeneously expressed in PDAC organoids. (A) Integrated UMAP clustering of PDAC organoids, annotated by cluster name (left) or organoid origin (right). (B) WNT expression in single cells of PDAC organoids. (C) UMAP with WNT grouping annotation. Cells expressing WNT7A, WNT7B or WNT10A (WNT^hi^) indicated in green, cells without expression of these WNTs (WNT^lo^) indicated in red. (D) Violin plots showing expression of indicated WNT target genes in WNT^hi^ and WNT^lo^ groups. (E) UMAP plots with subtype classification based on gene signatures from Raghavan *et al*., 2021. Top UMAP shows all cells, bottom UMAPs only display WNT^lo^ (left) and WNT^hi^ (right) cells. (F) Violin plot showing single cell subtype score of WNT^hi^ and WNT^lo^ groups (indicated Wilcoxon test p-value). (G) Volcano plot showing upregulated DEGs in WNT^+^ versus WNT^−^ groups. Selection of basal-like genes annotated in blue (Raghavan et al., 2021), FDR>0.05 threshold indicated with red dashed line. CLS: classical; BSL: basal-like; ns: not significant; DEGs: differentially expressed genes; IFN: interferon.

We next aimed to assess whether WNT secretion is linked to a specific cluster of PDAC cells. In line with bulk RNA sequencing results, we identified WNT7B, WNT10A and WNT7A as the most prominently expressed WNTs in single cell RNA sequencing data (Figure 4B, Figure S5B). Unexpectedly, WNT-secreting cells were not enriched within one specific cluster but were found scattered throughout all identified cell clusters (Figure 4C). Furthermore, most WNT-producing cells appeared dedicated to express a single WNT gene (Figure S4F). Because we cannot exclude the possibility that cells with very low WNT expression levels may have gone undetected, we refer to the two cell populations as WNT-high (WNT^hi^) and WNT-low (WNT^lo^). A comparison of gene sets between WNT^hi^ and WNT^lo^ cells revealed selective enrichment of genes involved in DNA replication in WNT^hi^ cells, while expression of WNT/β-catenin target genes were unaltered between both populations (Figure 4D, S4G). These findings thus suggest that although epithelial WNTs are expressed in a subset of WNT^hi^ PDAC cells, neighboring WNT^lo^ PDAC cells still display signs of active WNT signaling.

Subsequently, we tested whether the WNT-regulated signatures derived from LGK-treated PDAC organoids (Figure 3) are enriched in specific cell types of our single cell dataset. While WNT-driven genes were mainly expressed in Cycling-S and Diff1 clusters, genes usually inhibited by WNT were expressed by cells within the Diff2 population (Figure S4H). These data indicate that WNTs predominantly act by maintaining a cycling population, while preventing the transition towards Diff2 cells.

Recent reports using single cell PDAC tissue analysis suggested that PDAC classification in classical and basal subtypes is not binary, but rather reflects a gradual transcriptional state and, moreover, cells expressing markers of the two subtypes can reside in a single tumor (Krieger *et al*., 2021; Raghavan *et al*., 2021; Williams *et al*., 2023). We therefore examined subtype heterogeneity and their correlation with epithelial WNT expression levels within our PDAC organoid models. Indeed, we identified a heterogeneous distribution of subtypes at the single cell level within each PDAC organoid (Figure 4E, Figure S6). Notably, WNT^hi^ cells were clearly linked to a basal-like score in comparison to WNT^lo^ cells (Figure 4E,F). In addition, multiple genes that are part of the basal-like signature were significantly upregulated in WNT^hi^ cells compared to WNT^lo^ cells, including KRT7, UCA1 and CAV2 (Figure 4G). Taken together, PDAC organoids show intra-tumor subtype heterogeneity driven by a cell-intrinsic mechanism that correlates with WNT expression.

Subsequently, we assessed expression of previously published WNT-associated signatures, derived from human patient-derived PDAC organoids that either depend on an external source of WNT for growth (‘WNT-dependent’) or do not require WNT supplementation (‘WNT self-sufficient’) (Seino *et al*., 2018). Strikingly, both signatures displayed heterogeneous expression at the single-cell level within our PDAC organoid models, showing selective enrichment of the WNT-dependent signature within WNT^lo^ cell populations, while the WNT self-sufficient signature was enriched within WNT^hi^ cells (Figure S3I). These findings thus indicate that WNT-dependency states are highly divergent within PDAC tumors and are linked to the capacity of cells to secrete WNT.

### WNT7B reporter identifies basal-like cells in live organoids

To perform a comprehensive comparison of WNT^hi^ and WNT^lo^ PDAC cells, we designed a reporter for the identification of endogenous WNT7B-expressing cells by introducing an internal ribosome entry site (IRES) followed by a nuclear localization signal (NLS)-containing mNeonGreen (mNG) fluorophore, at a location downstream of the WNT7B locus (Figure 5A, Figure S7A). After integration, all reporter organoid lines revealed heterogeneous expression of mNeonGreen (Figure 5B), confirming the co-existence of cells with high (WNT7B^hi^) or low/undetectable WNT7B (WNT7B^lo^) expression within each PDAC organoid. Furthermore, P28 organoids contain a smaller fraction of WNT7B^hi^ cells in comparison to P40 and P47 organoids (Figure 5C).

**Figure 5.**
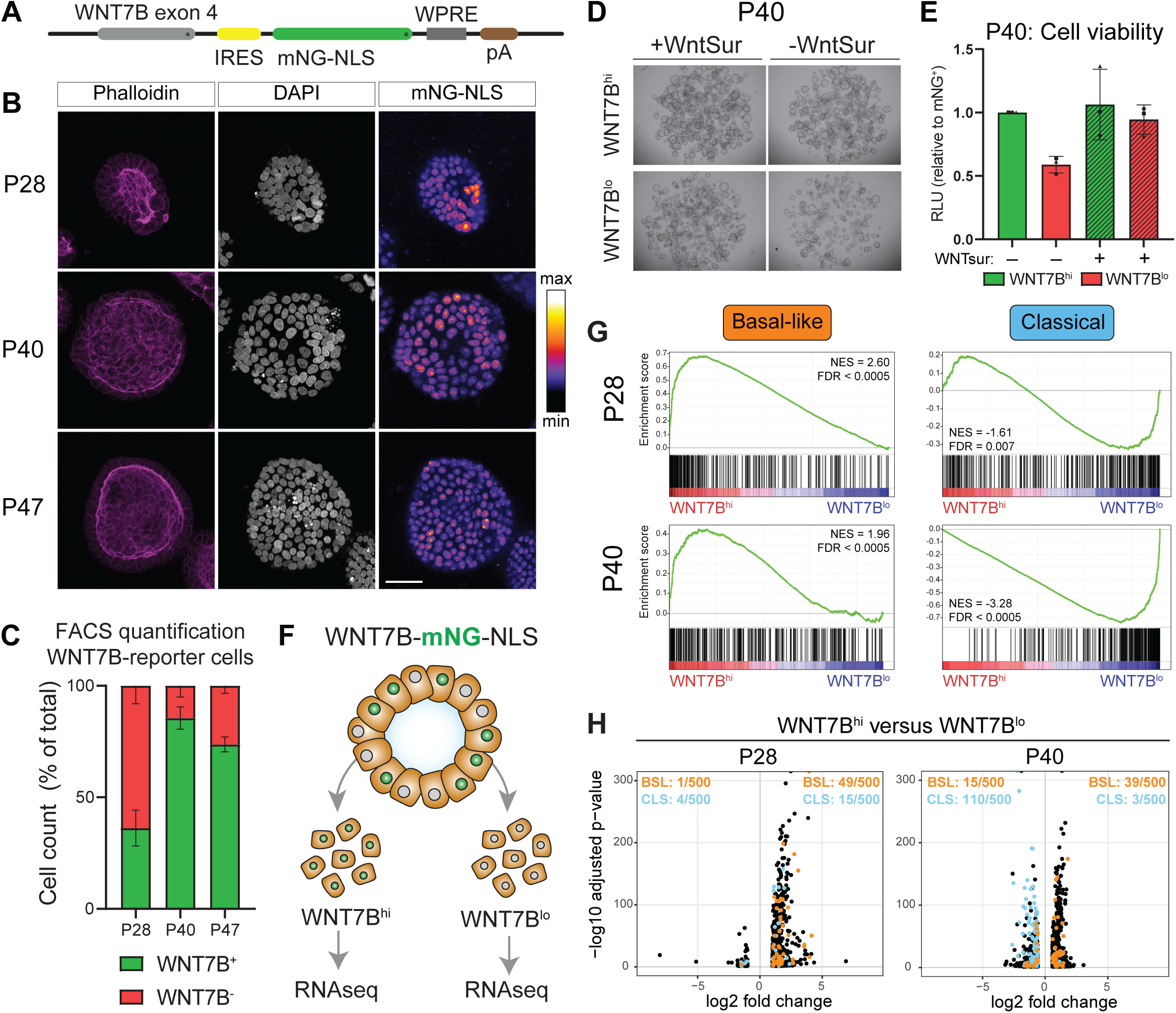
WNT7B reporter identifies basal-like cells in live organoids. (A) Schematic overview of integrated reporter construct. (B) Confocal microscopy maximum projection images of reporter organoids. Reporter signal visualized as Fire-lookup table. (C) FACS quantification of WNT7B^hi^ and WNT7B^lo^ cells, represented as percentage of total cells. (D) Brightfield images showing outgrowth of sorted WNT7B^hi^ and WNT7B^lo^ cells from P40, in absence or presence of WNTsur. (E) Cell viability assay of outgrowth experiment shown in (D). (F) Schematic overview of sequencing experiment on sorted cells. (G) GSEA of PDAC subtype signatures from Raghavan *et al*., 2021, comparing WNT7B^hi^ versus WNT7B^lo^ cells. (H) Volcano plot of DEGs in WNT7B^hi^ versus WNT7B^lo^ analysis. Annotated genes are present in top 500 basal-like (orange) and classical (blue) signature genes from Raghavan et al., 2021. Number of signature DEGs in WNT7B^hi^ and WNT7B^lo^ populations are displayed in top right and left, respectively, of each plot. IRES: internal ribosome entry side; WPRE: woodchuck hepatitis virus post-transcriptional regulatory element; NLS: nuclear localization signal. CLS: classical; BSL: basal-like. SB = 50µm.

To validate our reporter, we sorted WNT7B^hi^ and WNT7B^lo^ cell populations and assessed WNT7B expression using RT-qPCR. As expected, WNT7B^hi^ cells displayed higher expression of WNT7B in comparison to WNT7B^lo^ cells, thus confirming a correlation of mNG signal with endogenous WNT7B expression (Figure S7B). Next, we investigated the capacity of WNT7B^hi^ and WNT7B^lo^ cells to reconstitute organoid growth by sorting both cell populations from P28 and P40 reporter organoids (Figure S7C). Single WNT7B^hi^ P40 cells that were plated in the absence of exogenous WNT displayed more efficient outgrowth and organoid reconstitution compared to their WNT7B^lo^ counterparts (Figure 5D-E). The outgrowth of WNT7B^lo^ cells was fully rescued by supplementation with an exogenous source of WNT (Figure 5D-E), confirming that WNT deficiency underlies the impaired outgrowth of these cells . In line with the multi-WNT profile of P28, WNT7B^lo^ cells of this line were capable of growing out with a similar efficiency as WNT7B^hi^ cells (Figure S7D-E). Thus, WNT7B alone or along with other WNTs is essential for cell survival and proliferation in PDAC cells.

To perform an in-depth transcriptional profiling of WNT7B^hi^ and WNT7B^lo^ cells, we sorted the 25% highest and 25% lowest mNG-reporter cells and performed bulk RNAseq analysis of these populations (Figure 5F, Figure S8A-C, Table S3). GO-term analysis of overlapping DEGs in both P28 and P40 lines confirmed enrichment of cell cycle-related gene sets in WNT7B^hi^ cells (Figure S8D). In addition, WNT7B^hi^ cells were clearly enriched for expression of basal-like gene signatures, while WNT7B^lo^ cells were linked to transcriptional profiles of the classical PDAC subtype (Figure 5G,H; Figure S8E). Of note, in line with this observation, WNT7B^lo^ cells express commonly used classical-like PDAC tumor markers HNF1A and GATA6 (Figure S8F) that previously inversely correlated with WNT expression in PDAC samples (Brunton *et al*., 2020; Driehuis *et al*., 2019; Seino *et al*., 2018). In line with our single cell dataset (Figure S4I), WNT-dependency and WNT-self-sufficiency signatures (Seino *et al*., 2018) were enriched in WNT7B^hi^ and WNT7B^lo^ cells, respectively (Figure S8G). Taken together, our findings reveal that WNT7B^hi^ expression is a marker for a basal-like PDAC subtype state in WNT-dependent PDAC models.

### Endogenous WNT7B reporter reveals stable co-existence of cellular WNT states in live organoids

We wondered whether WNT7B expression is stable or dynamically expressed over time within PDAC organoids. To assess this issue, we performed timelapse imaging of reporter organoids with an integrated H2B-mCherry reporter to track nuclei over time. We processed the obtained timelapse images using an updated version of recently published organoid tracking software (Kok *et al*., 2020) and projected lineage trees of growing organoids. We observed that WNT7B reporter intensity remained stable in single cells over multiple cell divisions (Figure 6A). Although nuclear reporter levels dropped temporarily during cell division, cells recovered rapidly to reach similar reporter levels as before mitosis, thus stably maintaining high and low reporter cell lineages within the same organoid (Figure 6B). These data indicate stable co-existence of both WNT7B^hi^ and WNT7B^lo^ cell populations within PDAC organoids over time.

**Figure 6.**
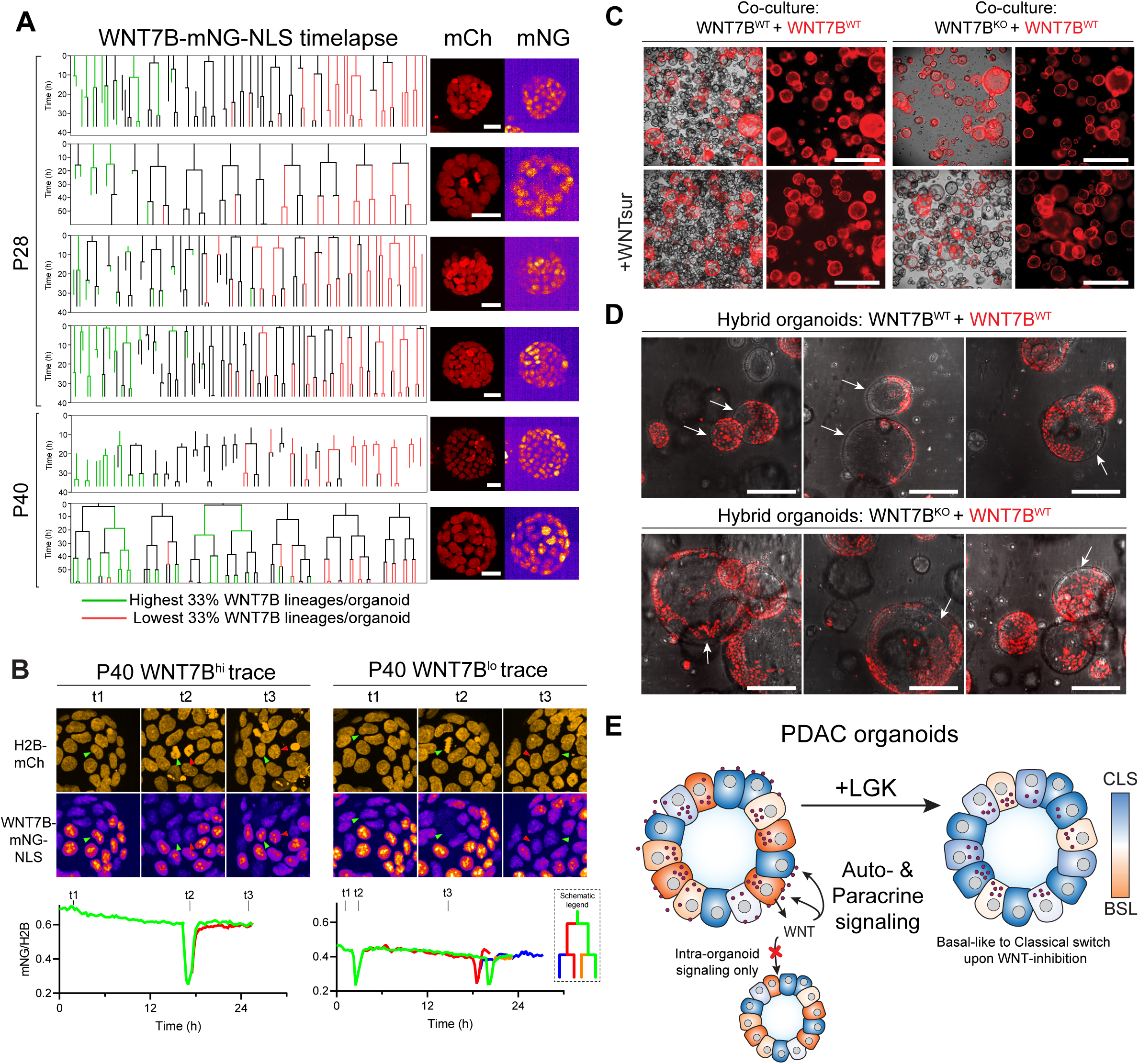
Endogenous WNT7B reporter reveals stable co-existence of cellular WNT states in live organoids. (A) Lineage trees of P28 and P40 organoids. Reporter signal is corrected over H2B-mCherry signal for each nucleus and then annotated as highest 33% (green) or lowest 33% (red) for each organoid. Maximum projection images of last frame of mCherry (mCh) and mNG channels displayed per lineage tree. (B) Still images (top) and reporter signal quantification (bottom) of timelapse imaging on P40 reporter organoids, showing signal stability before and after a cell division in in WNT7B^hi^ (left) and WNT7B^lo^ cell lineage. Single traces of indicated cells are shown. Scale bar = 25µm (C) Co-culture of WNT7B^WT^ H2B-mCherry P40 organoids with unlabeled WNT7B^KO^ (top) or WNT7B^WT^ (bottom) P40 organoid clones. Merged images of brightfield and mCherry signal displayed. Scale bar = 1000µm (D) Maximum projection images of hybrid culture from same organoid lines as in (C). Scale bar = 100µm (E) Schematic model. mCh: mCherry; mNG: mNeonGreen.

To assess interdependence of WNT7B^hi^ and WNT7B^lo^ populations within PDAC organoids, we examined the ability of WNT7B-expressing cells to rescue growth of WNT7B-deprived cells using our clonal P40 WNT7B^KO^ line. Upon co-culturing both organoid lines adjacent to each other, WNT7B^KO^ organoids failed to grow out, while WNT7B^WT^ cells were well-capable of reconstituting organoids growth in these conditions (Figure 6C), suggesting that WNT7B does not diffuse to neighboring organoids. Next, we generated hybrid cultures, in which single organoids are composed of both WNT7B^WT^ and WNT7B^KO^ cells (Garcia *et al*., 2021). Strikingly, within these hybrid organoids the outgrowth of WNT7B^KO^ cells was well-supported by neighboring WNT7B^WT^ cells, as shown by the formation of large patches of WNT7B^KO^ cells per organoid (Figure 6D). Based on these results, we conclude that WNT7B-expressing PDAC cells mediate local, short-range signaling to promote the survival and outgrowth of WNT-negative PDAC cells within the tumor tissue. Moreover, our findings explain how distinct WNT-linked cellular states can stably co-exist within WNT-dependent PDAC tumors.

## Discussion

WNT7B and WNT10A expression occurs in aggressive PDAC tumors and correlates with lower survival probability, but our understanding of how these specific WNTs drive pancreatic cancer progression remains incomplete. In this study, we show that (1) a subset of PDAC tumors employ intrinsic epithelial-derived WNT7B and WNT10A to drive WNT/β-catenin-mediated cancer cell growth and survival, (2) WNT-high PDAC cells are heterogeneously distributed within tumors and co-exist with WNT-low/negative PDAC lineages, (3) WNT-high cells drive survival and growth of neighboring WNT-low/negative cells via short range, cell-cell contact-dependent signaling, and (4) WNT7B maintains a more aggressive, basal-like PDAC subtype via suppression of a more differentiated, classical PDAC signature, and that this balance is shifted upon WNT inhibition (Figure 6E).

Over the past decade, multiple independent reports defined classical and basal-like transcriptional signatures as consensus PDAC subtype states (Bailey *et al*., 2016; Collisson *et al*., 2019; Moffitt *et al*., 2015). A growing body of evidence however suggests that PDAC cells occupy gradual rather than binary subtype states, and thus PDAC tumors are often comprised of hybrid cells that express both signatures simultaneously (Juiz *et al*., 2020; Raghavan *et al*., 2021; Saillard *et al*., 2023; Werba *et al*., 2023; Williams *et al*., 2023). Such tumor cell heterogeneity exists both in patient-derived PDAC biopsies (Chan-Seng-Yue *et al*., 2020; Raghavan *et al*., 2021; Werba *et al*., 2023) as well as *in vitro* in PDAC organoid models (Juiz *et al*., 2020; Krieger *et al*., 2021). Our data confirm PDAC subtype heterogeneity at the single cell level. We reveal that both WNT7B and WNT10A are heterogeneously expressed in PDAC tumors, and that expression of these WNTs correlates with a basal-like state. Moreover, we find that cellular heterogeneity is established and maintained by WNTs, via both paracrine and autocrine signaling. Indeed, perturbation of WNT signaling leads to loss of basal-like cells and thereby loss of heterogeneity.

Various studies indicate that microenvironmental signals and epigenetic changes dictate PDAC tumors to adopt a classical or basal-like subtype, and that these signals can rewire gene expression to mediate a transcriptional switch from one subtype to the other. A shift towards a basal-like state is mediated by either *GLI2* activation (Adams *et al*., 2019), *HNF4A* or *GATA6* loss (Brunton *et al*., 2020), TGFβ signaling (Raghavan *et al*., 2021), activity of transcription factor *TP63* (Somerville *et al*., 2018) or increased expression of splicing factor *Quaking* (Ruta *et al*., 2024). In addition, macrophages in the tumor microenvironment induce a basal-like state via TNFα/c-Jun signaling, while depletion of macrophage numbers via CSF1R inhibition induces a switch towards the classical subtype state (Candido *et al*., 2018; Tu *et al*., 2021). We complement these reports by showing that epithelial-intrinsic signaling via WNT7B and WNT10A suppresses the classical subtype state. Conversely, WNT inhibition shifts the transcriptional program towards a more classical state.

Since patients with classical-characterized tumors respond better to standard line chemotherapy and have longer overall survival rates (Aung *et al*., 2018; Suurmeijer *et al*., 2022), clinical perturbation of WNT7B/10A signaling to induce a basal-to-classical subtype switch might be of interest for combination therapy for WNT-dependent tumors. In addition, many PDAC tumors annotated as classical have a subpopulation of basal-like cells that enhances tumor aggressiveness and lowers overall survival probability compared to fully classical tumors (Saillard *et al*., 2023; Williams *et al*., 2023), suggesting a potential benefit from a WNT inhibition-induced subtype shift that yields a more homogenous classical PDAC tumors. In line with these observations, patient-derived organoid models showed that chemosensitivity is coupled to the transcriptional subtype state (Krieger *et al*., 2021; Raghavan *et al*., 2021).

Although detailed subtyping of PDAC cells relies on transcriptomic analysis, several surrogate markers such as GATA6 and TFF1 are used to identify classical tumor cells, while KRT5 and S100A2 mark basal-like regions (O’Kane *et al*., 2020; Williams *et al*., 2023). Our results argue that WNT7B expression may be employed as an alternative basal-like subtype surrogate marker. Furthermore, existing markers used for diagnostic purposes are visualized by conventional staining procedures on formalin-fixed paraffin-embedded tissue and therefore unable to identify subtype plasticity in live tumor cells. We show that our endogenous WNT7B reporter can function as a live marker for the basal-like subtype state in WNT-dependent PDAC organoids. With this model system, we open up the possibility of using live imaging to assess alterations in PDAC subtype plasticity during conventional chemotherapy treatment as well as potential novel targeted therapies, in both *in vitro* and *in vivo* PDAC tumor models.

The pathways downstream of WNT7B and WNT10A during healthy cellular and developmental processes have remained incompletely understood. Multiple studies suggest that WNT7B and WNT10A can drive canonical (Alok *et al*., 2017; Arensman *et al*., 2014; Stenman *et al*., 2008) and β-catenin-independent, non-canonical WNT signaling (Kimura *et al*., 2020; Zheng *et al*., 2013; Zhong *et al*., 2022) in various organ types and physiological states. Our transcriptional analysis of LGK-treated PDAC organoids clearly revealed that WNT7B and WNT10A drive β-catenin-dependent WNT target gene expression. Recently, Zhong and colleagues reasoned that epithelial-derived WNTs (WNT7A/B, WNT10A) drive non-canonical WNT signaling, while mesenchymal WNTs (WNT2/2B) activate a β-catenin dependent program (Zhong *et al*., 2022). Since our *in vitro* models lack a mesenchymal component, our results convincingly show that WNT7B and WNT10A sustain intrinsic WNT/β-catenin signaling and drive growth of the PDAC cancer epithelium, even when expressed only in a subpopulation of cells.

While many research efforts have focused on studying cancer stem cells, the possible role of cancer stem cell-supporting niche cells has been relatively unexplored. Although non-epithelial cells, such as fibroblasts (Hutton *et al*., 2021) and endothelial cells (Choi *et al*., 2021), can support cancer cells, the presence of functional niches within the tumor epithelium itself has been poorly explored. Specialized cancer niche cells that promote cancer stem cell growth by providing WNT ligands were described for only a few cancer models, including lung adenocarcinoma, with a prominent role for WNT7B- and WNT10A-secreting niche cells (Tammela *et al*., 2017) and colorectal cancer, that harbor tumor intrinsic niche cells via RNF43 mutations (Bugter *et al*., bioRxiv, 2023). These cancer niche cells are characterized by features of differentiation and expression of several niche factors, including IGF2, DLL1/4 and various WNTs (Tammela *et al*., 2017; Bugter *et al*., bioRxiv, 2023). Our results indicate that PDAC make use of a different WNT-driven growth model. First, both WNT-positive and WNT-negative cell populations display expression of a WNT/β-catenin transcriptional program and both populations actively undergo cell division. Second, WNT7B-expressing cells display features of both autocrine and paracrine signaling. Third, we show that WNT signaling occurs at short range and requires cell-cell contact, in line with earlier reports (Farin *et al*., 2016; Kimura *et al*., 2020; Zheng *et al*., 2013). We postulate that our findings are analogous to the role of WNT7B and WNT10A in tissue regeneration, where cells are induced to (re-)enter the cell cycle and contribute to wound healing and tissue repair.

The mechanism by which PDAC tumors drive WNT7B and WNT10A expression remains unclear. Recent work indicates that WNT self-sufficiency in gastric cancer may arise either through copy number variation or through the acquirement of KRAS mutations (Teriyapirom *et al*., bioRxiv, 2023), events that also frequently occur in PDAC tumors (Chan-Seng-Yue *et al*., 2020). We observe no specific gain of the WNT7B locus in DNA sequencing analysis of PDAC organoids, suggesting alternative routes to increased WNT expression levels. Other signaling pathways such as YAP (Volckaert *et al*., 2017) and TGFβ (Oda *et al*., 2016; Sundqvist *et al*., 2020) converge on the WNT signaling axis by driving WNT7B expression. Furthermore, epigenetic alterations can result in WNT pathway deregulation, including WNT7B and WNT10A expression in basal-like PDAC (Brunton *et al*., 2020; Nicolle *et al*., 2017). On the other hand, GATA6 was shown to negatively regulate WNT expression, as GATA6 inactivation upregulates WNT7B and renders organoids independent of WNT supplementation to culture medium (Seino *et al*., 2018). A more detailed understanding of regulation of WNT ligand expression in cancer context would be an interesting topic for future investigation.

WNT-addicted pancreatic cancer models, including cell-line based (Steinhart *et al*., 2016), xenograft (Zhong *et al*., 2019) and organoid (Seino *et al*., 2018) models, are sensitive to PORCN-inhibition that shows synergistic effects with PI3K/mTOR-pathway inhibition (Zhong *et al*., 2019) and GSK3β-inhibition (Brunton *et al*., 2020). PORCN inhibitors however are not yet successful in the clinic, partly due to their side effects (Ng *et al*., 2017; Rodon *et al*., 2021). RNF43 mutations predict PORCN-inhibitor sensitivity *in vitro* (Jiang *et al*., 2013) and is included as a selection criterion for patient inclusion in clinical trials involving WNT perturbation (NCT01351103). Our data however suggest that RNF43-WT patients who express epithelial WNTs might also benefit from pharmacological WNT inhibition. Alternatively, improved stratification of patients may be required to better predict efficacy of pharmacological WNT perturbation, as PDAC classification based on WNT dependency revealed that some PDAC tumors are WNT- and RSPO-independent, and this state is linked to p300 status (Seino *et al*., 2018; Zhong *et al*., 2022). Our data show that a subset of PDAC patients with WNT-dependent, basal-like characteristics may benefit from WNT7B-specific inhibition, either alone or in combination with chemotherapy, which is likely to be accompanied with reduced side-effect toxicity compared to broad PORCN-mediated WNT inhibition that systemically affects all WNT niches and WNT-dependent tissues (Werner *et al*., 2023).

In summary, our results shed light on how cancer cell-derived WNT7B and WNT10A promote PDAC growth and sustain cancer cell heterogeneity and suggest that interference with a specific subset of WNTs may provide a therapeutical strategy to force tumor class switching to a less aggressive cancer subtype that correlates with improved clinical responses.

## Methods

### Patient material and data

The collection of pancreatic tissue for the generation patient-derived organoids was performed according to the guidelines of the European Network of Research Ethics Committees (EUREC) following European, national and local law. All donors participating in this study provided signed informed consent. PDAC organoid lines with identified codes HUB-08-B2-028A, HUB-08-A2-028C, HUB-08-B2-040A, HUB-08-A2-040C, HUB-08-B2-047A and HUB-08-A2-047C are cataloged by Hubrecht Organoid Technology (https://www.huborganoids.nl/) and can be requested at. The Biobank Research Ethics Committee of University Medical Center Utrecht (TCBio) approved the release protocol (TCBio number 19-188) under which this research was performed. Distribution to third (academic or commercial) parties will have to be authorized by the Biobank Research Ethics Committee of the University Medical Center Utrecht (TCBio) at request of Hubrecht Organoid Technology.The SPACIOUS cohort was interrogated for PDAC patient RNA expression data (Dijk *et al*., 2020; Suurmeijer *et al*., 2022).

### Organoid culturing

PDAC organoids were seeded in drops of Matrigel (Corning) or Basement Membrane Extract (BME, R&D systems) and passaged every 7-10 days (P28 and P40) or every 8-12 days (P47) using mechanical disruption and seeded into a 1:3 to 1:4 ratio. Matrigel/BME was diluted in base medium (see below) to 70%. For experiments, organoids were made into a single cell expansion using TrypLE Express (Life Technologies) incubation at 37 °C and filtered through a 40µm strainer before seeding. Organoids were established and cultured in PDAC tumor medium (Driehuis *et al*., 2019) consisting of advanced DMEM/F12 medium (Invitrogen), 1% Penicillin/Streptomycin (P/S, Lonza), 1% HEPES buffer (Invitrogen), 1% Glutamax (Invitrogen), referred to as base medium, which was further supplemented with 10% RSPO-CM (in-house), 2% Noggin-CM (in-house), 1x B27 (Invitrogen), 1.25 mM N-acetylcysteine (Sigma-Aldrich), 10 mM Nicotinamide (Sigma-Aldrich), 100 ng/mL human recombinant FGF10 (Peprotech), 50 µg/mL Primocin (InvivoGen), 5 ng/mL human recombinant EGF (PeproTech) and 10nM Gastrin (Sigma-Aldrich), referred to as PDAC medium. Culture medium was refreshed every 2-3 days. For inhibition of WNT secretion, organoid culture medium was supplemented with 200nM LGK-974 (Selleckchem). Where indicated, organoid medium was supplemented with 50% WNT3A conditioned medium (WNT3A-CM, in-house), L-cell conditioned medium (LCM, in-house) or a conditioned medium-variant of WNT-surrogate molecule (Janda *et al*., 2017) at a concentration of 0.5-1% (WNTsurCM, in-house). WNTsurCM was obtained by harvesting and filtering conditioned medium of PEI-transfected HEK293T cells after 4 days. WNTsurCM activity was validated using TCF/LEF-reporter assay, measured with Dual Luciferase Reporter Kit (Promega) and SDS-PAGE Western blot analysis. WNTsurCM concentration was adjusted for each batch.

### Plasmids and CRISPR design

WNT knock-out organoid lines were generated using pSpCas9(BB)-2A-Puro (Addgene, 48139) backbone plasmids, in which the following guide RNAs were integrated using golden gate cloning: WNT7B guide: CGCGGTGATGGCGTACGTGA; WNT10A guide: TGAAGATGGGACTCTCATAG. WNT7B-reporter knock-in were generated using pSpCas9n(BB)-2A-Puro (Addgene, 62987), in which the following guide RNA was introduced by golden gate cloning: CATAGACGGGTGCAGAAGCG. Q5 High Fidelity 2X Mastermix (NEB) and In-Fusion HD Cloning kit (Takara) used for general cloning procedures. The targeting vector (TV) was generated using a SapI golden gate cloning method (Bollen *et al*., 2022), inserting homology arms of ∼600bp (DNA fragments, Genscript).

### Lentiviral transduction

Lentivirus containing an H2B-mCherry construct (pLV-EF1a-H2B-mCherry-IRES-blast; kind gift from the S.M.A. Lens lab) was generated in HEK293T cells according to standard procedures and concentrated using Lenti-X^TM^ Concentrator (Takara, 631231). Viral incubation performed by combining a single cell suspension of PDAC organoids with concentrated virus for 4h in PDAC medium without primocin and P/S, supplemented with 8µg/ml Polybrene (Merck) and 10 µM Y-27632 (Selleck chemicals). Selection of H2B-tagged lines was done using 5µg/ml blasticidin (InvivoGen).

### Electroporation

Organoids were made single cell using TrypLE Express (Life Technologies) at 37°C. Electroporation (EP) was performed using NEPA21 (NEPAGENE) electroporator, following previously described methods for human organoids (Fujii *et al*., 2015). For CRISPR-mediated knock-out experiments, 4µg of WNT-targeting CRISPR plasmids, 3.6ug of pPB-CAG-rtTA_Hygromycin and 2.6ug of PiggyBac Transposase was introduced. For generation of knock-in lines, 12.5ug of TV and 7.5ug WNT7B-targeting CRISPR plasmid. After electroporation, organoids were cultured in PDAC culture medium without P/S and primocin for at least 4 days, and in presence of 1.25% DMSO for 1 day and 10 µM Y-27632 (Selleck chemicals) for at least 2 days while outgrowth of cells was monitored. For specific WNT knockout experiments, medium was supplemented with 0.5-1% Wnt surrogate conditioned medium (WNTsurCM, in-house) when indicated. Selection was performed using 2µg/ml puromycin (Sigma-Aldrich) or 100 µg/ml hygromycin (Merck). Clonal lines were generated by manually isolating organoids and subsequently expanding them. Selection cassette (EF1A-Ruby-T2A-Puro) from reporter organoids was removed by electroporating 10µg of a transiently expressed Cre-lox plasmid (Addgene, #55632; kind gift from the H.J.G. Snippert lab).

### Immunofluorescence and microscopy

Organoids were seeded in BME in a 15-well Ibidi chamber (IBIDI) after passaging and fixed 5 days later Fixation was done in 4% Paraformaldehyde (Sigma-Aldrich), diluted in 0.1M phosphate buffer for 30 minutes at room temperature (RT). After fixation organoids were washed and permeabilized with PBD0.2T (0.2% Triton X-100, 1% DMSO and 1% Bovine Serum Albumin (BSA; Roche) in PBS), after which they were incubated with primary antibodies diluted in PBD0.2T buffer at 4°C overnight. After washing in PBD0.2T, secondary antibodies were added the next day and incubated 2-3 hours at RT. Primary antibodies and dyes used are anti-MUC5AC (1:500; ThermoFisher, MA5-12178), Goat-anti-mouse Alexa Fluor^TM^ 568 (1:200; Invitrogen, A-11031), DAPI (1:1.000; Sigma, D9542) and Phalloidin-Alexa-647 (1:200; CST, 8940S). Stained organoids were stored and imaged in IBIDI mounting medium (IBIDI). Images were acquired using a confocal Zeiss LSM880 microscope. Brightfield images of organoids in culture were taken using an EVOS imaging system (Thermo Fisher Scientific).

### Timelapse imaging

Reporter organoids from bulk populations of P28 and P40 organoids were either dissociated into single cells using TrypLE Express (Life Technologies) and filtered through a 40 µm nylon cell strainer (Falcon) or split mechanically into small fragments before seeding in an 8-well IBIDI imaging slide (IBIDI). Timelaps imaging started at day 2 or day 4 post seeding and performed for either 37h or 60h. Imaging performed on a Nikon Ti2-E stand with Perfect Focus System (PFS) and equipped with the Yokogawa CSU-W1 Spinning Disk Confocal Unit and 2 scientific CMOS camera’s from Andor (Zyla 4.2 Plus). The objective used was a 20x CFI Plan Apochromat Lambda Air, NA 0.5, WD 1.0mm. The system is controlled using the Nikon NIS Elements AR Software (v5.20).

Single trace analysis was performed using Trackmate (Ershov *et al*., bioRxiv, 2021) and StarDist (Schmidt *et al*., 2018) plugins in ImageJ, quantifying max-intensity projections with single nucleus reporter signal correction over corresponding H2B-mCherry signal.

Whole-organoid lineage tracking was performed using OrganoidTracker (Kok *et al*., 2020). Fluorescence signal was measured in 3D within a sphere positioned at the nucleus center with a radius of 3 µm and normalized by dividing with the H2B-mCherry signal of each nucleus. For every cell trajectory, the average WNT signal was calculated in between 2 mitotic events by averaging the signal across all time points. Short cell trajectories (spanning less than 33% of the time lapse duration) that move out of the field of view were filtered out for further analysis. Lineage trees were plotted using a custom-written Python script, which orders all lineages based on their average WNT signal.

### Hybrid organoid generation and co-culture

Hybrid organoids were generated using an adapted protocol from (Garcia *et al*., 2021). Briefly, clonal H2B-mCherry tagged organoids (cultured in PDAC organoid medium) and clonal WNT7B^WT^ or WNT7B^KO^ organoids were both cultured in PDAC organoid medium supplemented with 0.5% WNTsurCM. Two days prior to the experiment, medium was exchanged for PDAC organoid medium without WNTsurCM. On day 0, all organoids were dissociated and strained into single cells using TrypLE Express (Life Technologies), and subsequently pre-mixed in equal amounts in 30µl PDAC medium. After a centrifugation step, cell pellets were incubated at 37°C for 1 hour, followed by plating organoids in BME in an 8-well ibidi imaging plate (IBIDI). Organoids were imaged at day 7 after plating on a Nikon Spinning Disk Confocal Unit (see above).

For co-culture experiments, the same organoid lines and single cell dissociation procedure was followed as for hybrid organoid generation, but pre-mixed single cell suspensions were not incubated before plating and seeded directly in BME. Images were taken at 7-9 days after seeding on a Leica Thunder microscope, using a 5x objective HC PL Fluotar, NA 0.15, WD 13.7.

### RNA isolation and RT-qPCR

A total of 100-150µl of Matrigel drops with organoids was harvested for RNA isolation using RNeasy Mini Kit (Qiagen) according to manufacturer’s protocol. Organoids were cultured and passaged once after fluorescence-activated cell sorting (FACS) and harvested at day 9 after seeding using RLT lysis buffer (Qiagen) with 1% 2-mercaptoethanol. Turbo DNAse (Thermo Fisher Scientific) was used to remove genomic DNA. cDNA was synthesized using iScript cDNA synthesis kit (Bio-Rad) and subsequently used in qRT–PCR using IQ SYBR green mix (Biorad) according to the manufacturer’s protocol. Primers for WNT7B: CCCCCTCCCTGGATCATGCACA (forward) and GCCACCACGGATGACAGTGCT (reverse). ActinB primers were used as a housekeeping gene: GAAGCCGGCCTTGCACAT (forward) and AGCACAGAGCCTCGCCTTT (reverse).

### RNA sequencing

Three biological replicates of organoids were harvested at indicated timepoints (Figure 3) or sequenced directly after FACS (Figure 5F). RNA was obtained using the Qiagen QiaSymphony SP (Qiagen) according to the manufacturer’s protocol. Library prep was performed using Truseq RNA stranded polyA (Illumina). Library prep was performed using Truseq RNA stranded polyA (Illumina). Libraries were sequenced on an Illumina NextSeq2000 platform with 1×50bp reads. Quality control on the sequence reads from the raw FASTQ files was done with FastQC (v0.11.8). TrimGalore (v0.6.5) as used to trim reads based on quality and adapter presence after which FastQC was used to check the resulting quality. rRNA reads were filtered out using SortMeRNA (v4.3.3) after which the resulting reads were aligned to the human reference genome (GCA_000001405.15_GRCh38_no_alt_plus_hs38d1_analysis_set) using the STAR (v2.7.3a) aligner. Followup QC on the mapped (bam) files was done using Sambamba (v0.7.0), RSeQC (v3.0.1) and PreSeq (v2.0.3). Readcounts were then generated using the Subread FeatureCounts module (v2.0.0) with the GRCh38.106 gtf file as annotation. CPM and RPKM Normalized versions of the counts table were generated using the R-packages edgeR (v3.28) as well as a normalized version using DEseq2 (v1.28) (Love *et al*., 2014).

Expression data (RPKM) of PDAC and wildtype pancreas organoid lines (Figure S1C) was obtained by HUB Organoids (https://www.huborganoids.nl/).

### PurIST scoring

Bulk RNAseq data from SPACIOUS cohort (Dijk *et al*., 2020) and SPACIOUS-2 cohort (Suurmeijer *et al*., 2022) was analyzed and scored using PurIST classification (Rashid *et al*., 2020). For comparing gene expression on different PurIST classes, “lean classical” and “likely classical” groups as well as “lean basal” and “likely basal” were combined.

### FACS

Single cells from organoid were harvested using TrypLE incubation at 37 °C and filtered through a 40 µm nylon cell strainer (Falcon). Before sorting, cells were treated with DRAQ7 (1:1000; BIOKÉ) to gate on cell viability. FACS was performed on FACS Aria III machine (BD). Populations were gated for viable single cells and gated as indicated. Sorted cells were directly lysed in RLT lysis buffer (Qiagen) for RNAseq analysis (Figure 6) or plated in BME for organoid reconstitution (Figure 5D-E). FACS data were analyzed using the Cytobank platform (Kotecha *et al*., 2010).

### Single cell RNA sequencing

Variable single cells were FACS sorted into 384-well cell capture plated (Single Cell Discoveries). Each well of a cell capture plate contains a small 50nl droplet of barcoded primers and 10µl of mineral (Signa, M8410). After sorting, plates were immediately spun and placed on dry ice. Plated were stored at -80°C.

scRNA-seq was performed by Single Cell Discoveries according to an adapted version of the SORT-seq protocol (Muraro *et al*., 2016) with primers described in (Van Den Brink *et al*., 2017). Cells were heat-lysed at 65°C followed by cDNA synthesis. After second-strand cDNA synthesis, all the barcoded material from one plate was pooled into one library and amplified using *in vitro* transcription (IVT). Following amplification, library preparation was done following the CEL-Seq2 protocol (Hashimshony *et al*., 2016) to prepare a cDNA library for sequencing using TruSeq small RNA primers (Illumina). The DNA library was paired-end sequenced on an Illumina NExtseq Libraries were sequenced on Nextseq™ 500, high output, with a 1×75 bp Illumina kit_J(read 1: 26 cycles, index read: 6 cycles, read 2: 60 cycles). Reads were mapped to the hg38 reference genome. Analysis was performed with the R-package Seurat (4.3.0). Low quality cell barcodes having either less than 3000 unique transcripts, less than 1000 unique features or more than 50% mitochondrial transcripts were excluded. External RNA controls (ERCCS) and transcripts mapping to the mitochondrial genome were removed from all 908 cells that passed quality control. Data integration was performed following previously described method (Stuart *et al*., 2019). Cell cycle state was assessed by built in cell cycle signatures within Seurat. The first 17 principal components were used both to calculate dimensionality reduction using UMAP and to perform clustering with a resolution of 0.6. Scores for gene signature expression were calculated using the AddModuleScore function. Top 50 genes from WNT-signature from this paper (Figure 3) was used in Figure S4H. WNT groups were defined by indicated WNT gene expression being larger than 0 (WNT^hi^) or equal to 0 (WNT^lo^).

### Subtype scoring

Z-score of Raghavan subtype signatures on bulk RNA-seq data was calculated using GSVA analysis (see below), using the top 500 genes for each subtype. Subtype scoring on single cell data was performed as in (Krieger *et al*., 2021). Briefly, top 30 genes for classical and basal-like subtype signatures from (Raghavan *et al*., 2021) were used to calculate subtype scores for basal-like (S_bas_) and classical (S_cla_) for each individual cell using the AddModuleScore function in Seurat. Combined subtype score (S_sub_= max[0, S_cla_] − max[0, S_bas_]) was calculated for each cell. For WNT dependency scores of (Seino *et al*., 2018) on for single cell data, WNT10A gene was removed from WNT self-sufficient signature to not bias WNT^hi^ and WNT^lo^ grouping.

### DNA sequencing

FASTQ files of tumor DNA organoid data were used as input for the standard Hartwig analysis workflow. The Hartwig somatic calling pipeline (https://github.com/hartwigmedical/pipeline5) is hosted on a Google Cloud platform using platinum https://github.com/hartwigmedical/platinum that enables to run the complete Hartwig pipeline at once. A full pipeline description is explained in (Priestley *et al*., 2019), and details and settings of all the tools can be found on their Github page (https://github.com/hartwigmedical/hmftools). Briefly, reads were mapped to GRCh38 using BWA (v0.7.17). GATK (v3.8.0) Haplotype Caller was used for calling germline variants in the reference sample. SAGE (v2.2) was used to call somatic single and multi nucleotide variants as well as indels. GRIDSS (v2.9.3) was used to call simple and complex structural variants. PURPLE combines B-allele frequency (BAF) from AMBER (v3.3), read depth ratios from COBALT (v1.7), and structural variants from GRIDSS to estimate copy number profiles, variant allele frequency (VAF) and variant clonality. Mutations listed in the PRUPLE driver catalog tables were used to identify the driver variants.

### Cell Viability Assay

Quantitative viability assays were performed by CellTiter-Glo (Promega) according to manufacturer’s instructions.

### Data Analysis

Gene set enrichment analysis (GSEA, 4.3.2) was performed using the Broad Institute GSEA tool (software.broadinstitute.org/gsea/index.jsp) with default settings, using permutation by gene sets. Image processing was performed using ImageJ (2.3.0). Gene set variation analysis (GSVA) was performed using GSVA package (1.42.0) in R (4.1.2). Plots were generated using R packages pheatmap (1.0.12), ComplexHeatmap (2.10.0) or ggplot2 (3.4.0), bar graphs were generated with Graphpad Prism 9. GO-term analysis were performed online (Aleksander *et al*., 2023). Venn diagrams were created using Venny (Oliveros, 2015). Benchling software used for sequencing alignment. Synthego ICE tool was used to determine knock-out scores (Conant *et al*., 2022).

### Statistics

Statistics performed in Graphpad Prism 9 or R (4.1.2). For single cell data: Shapiro-Wilk normality test performed on subtype scoring for WNT groups in separate lines. Only P28 had normally distributed data, for consistency a nonparametric Wilcoxon test was performed. For GSVA analysis on bulk RNA sequencing data in Figure 3H: Shapiro-Wilk test indicates normality for all conditions except the Raghavan_classical signature in P47 (p=0.0444). However, Kolmogorov–Smirnov test indicates normality for all conditions (p>0.0100). Therefore, t-test performed and significance is indicated in the plot.

## Data availability

The scRNA-seq and bulk RNA-seq data generated in this study will be uploaded to the European Genome-phenome Archive (EGA) upon publication. If needed prior publication, data can be requested via the corresponding author.

## Author contributions

J.S., J.Y.V. and M.M.M. conceived and designed experiments. J.S. and D.X. performed experiments. J.S. and R.N.U.K. performed imaging data analysis. J.S. and J.B. performed reporter construct design and generation. J.S., J.Y.V. and M.M.M. analyzed the data and wrote the manuscript, which was reviewed by all authors. F.D. and L.A.A.B. provided patient expression data.

## Acknowledgements

WNT surrogate plasmid was a kind gift from C.Y. Janda. Cre-lox plasmid was a kind gift from H.J.G. Snippert. H2B-mCherry lentiviral plasmid was a kind gift from S.M.A Lens. MUC5AC antibody was a kind gift from M. Gloerich.

