## Supplementary_figures&legends for "WNT7B drives a program for pancreatic cancer subtype switching and progression"

### Supplementary figure legends

**Figure S1 – WNT expression in PDAC tissue samples and organoids.**
(A) Heatmap showing expression of all WNTs in PDAC tissue samples of SPACIOUS cohort. (B) Expression of WNT7B, WNT7A, WNT10A and PORCN in subtype-annotated PDAC tissue samples. P-values of Kruskal-Wallis test indicated. (C) Expression (RPKM) of WNT7B, WNT10A and WNT7A in selected PDAC organoid lines and corresponding WT organoid lines from the same patient. (D) Brightfield imaged of selected PDAC organoid lines. (E) WNT expression in PDAC organoids, expressed as percentage of total WNT expression (DEseq2 normalized counts). Percentages above 3% are indicated. Scale bar = 500µm.

**Figure S2 – PDAC organoids depend on endogenous WNT proteins.**
(A) Brightfield images of outgrowth experiment with indicated rescue conditions (passage 0, day 7). (B) Schematic representation of intra- and inter-organoid WNT signaling in a bulk population of wildtype and WNT$^{KO}$ organoids. (C-D) DNA and protein sequences of WNT7B$^{KO}$ (C) and WNT10A$^{KO}$ (D) clonal P40 organoid lines. Reference wildtype DNA and protein sequences are indicated, mutant nonsense sequence is indicated in red. Guide sequences for Cas9-targeting are highlighted. (E-F) Genotyping results of WNT7B (E) and WNT10A (F) clonal knock-out organoid lines. (G) DNA and protein sequences of WNT7B/10A$^{DKO}$ clonal P28 organoid line. Mutant nonsense sequence is indicated in red. (H) Brightfield images of WNT7B/10A double knock-out P28 organoid clone outgrowth experiment in presence and absence of WNTsur. After passage 3, LGK-974 was added to the control condition test for WNT dependency of this WNT7B/10A$^{DKO}$ clone. SB = 1000µm.

**Figure S3 – WNT proteins drive PDAC cell proliferation and prevent classical subtype gene expression.**
(A) Brightfield images of PDAC organoid lines after 24h of DMSO- or LGK-treatment. (B-C) Heatmaps (B) and bar plots (C) of core WNT target gene expression in DMSO- and LGK974-treated PDAC organoids. (D) GSVA analysis of gastric metaplasia and damage signature (Chen et al., 2021). Asterisks indicate p-values (t-test, P28: p=3e-04, P40: p=0.0054, P47: p=0.0028). (E) GSEA analysis of Gastric mature pit cell gene signature (Busslinger et al., 2021) on all PDAC lines. NES: Normalized enrichment score; SB = 1000µm.

**Figure S4 – PDAC-derived WNTs are heterogeneously expressed in PDAC organoids.**
(A) UMAP of organoid lines without integration. (B) Quality control UMAP plots of integrated single cell sequencing data with annotated number of genes per cell (left) and read count per cell (right). (C) Cell cycle phase annotation of single cells. (D) Heatmap of top differentially expressed genes for each cluster. (E) Heatmap of top differentially expressed genes between clusters Diff1 and Diff2. (F) Barplot indicating number of cells (expressed as percentage of total cells) that express either a single WNT or co-express multiple WNTs. (G) GO term biological processes analysis on differentially expressed genes between WNT$^{hi}$ and WNT$^{lo}$ cells. Dashed line indicated FDR=0.05. (H) WNT core signatures from Figure 3 projected on integrated single cell data, as module score displayed in a violin plot. Lowly-expressed genes from bulk RNA sequencing results (<100 average expression) were not included in this analysis. (I) Violin plots of WNT grouped single cell

data, showing module scores of WNT (in)dependency signatures from Seino *et al*., 2018. Asterisks indicate significance levels (Wilcox test, WNT-self-sufficient signature P28: p=0.004, P40: p=0.0095; P47: p=0.78; WNT dependency signature P28: p=0.0078; P40: p=3.2e-05; P47: p=0.00087). IFN: interferon; ns: not significant.

**Figure S5 – Single cell sequencing cluster analysis.**
(A) GO biological processes analysis for each cluster. (B) Violin plots of integrated single cell data, showing WNT protein expression. WNTs with detectable expression in less than 5 cells or 5-10 cells with expression level below 0.5 were not shown. WNT of interest in this paper are highlighted in bold.

**Figure S6 – Separate organoid line Seurat analysis.**
(A) Separate organoid line Seurat analysis showing for each organoid line (from left to right): clusters, cell cycle phase, single cell subtype classification scores (based on Raghavan *et al*., 2021) and WNT grouping annotation. (B) Violin plots showing single cell subtype score of WNT$^{hi}$ and WNT$^{lo}$ groups. Asterisks indicate level of significance (Wilcoxon test, P28: p=0.0065; P40: p=3.77e-06; P47: p=4.2e-05). CLS: classical; BSL: basal-like.

**Figure S7 – WNT7B-reporter supplemental image.**
(A) Detailed schematic of reporter construct, showing untargeted WNT7B locus (top), integrated full reporter construct (middle) and integrated reporter construct after removal of floxed downstream selection cassette (bottom). (B) qPCR for WNT7B on sorted mNG$^{hi}$ and mNG$^{lo}$ cell populations. (C) FACS histograms showing gate settings from mNeonGreen on control (top) and reporter (bottom) P28 and P40 cells. (D) Brightfield images showing outgrowth of sorted WNT7B$^{hi}$ and WNT7B$^{lo}$ cells from P28, in absence or presence of WNTsur. (E) Cell viability assay of outgrowth experiment shown in (D). DHA: downstream homology arm; ins: insulator; EF: elongation factor-1 alpha promoter; IRES: internal ribosome entry side; NLS: nuclear localization signal; pA: poly-A signal; UHA: upstream homology arm; WPRE: woodchuck hepatitis virus post-transcriptional regulatory element.

**Figure S8 – Transcriptional analysis of WNT7B$^{hi}$ and WNT7B$^{lo}$ cells.**
(A) FACS histrogram plots showing gating settings for reporter populations. (B) WNT expression in sorted populations. (C) Venn diagrams showing overlap of up- and downregulated genes in WNT7B$^{hi}$ versus WNT7B$^{lo}$ cells in P28 and P40 organoids. (D) GO-term analysis of overlapping DEGs in WNT7B$^{hi}$ cells in both P28 and P40 organoids. (E) GSEA plots of Moffit *et al*., 2014 signatures. (F) Heatmap of classical regulatory genes in P28 and P40. (G) GSEA plots of WNT (in)dependency signatures (Seino et al., 2018).

Figure S1

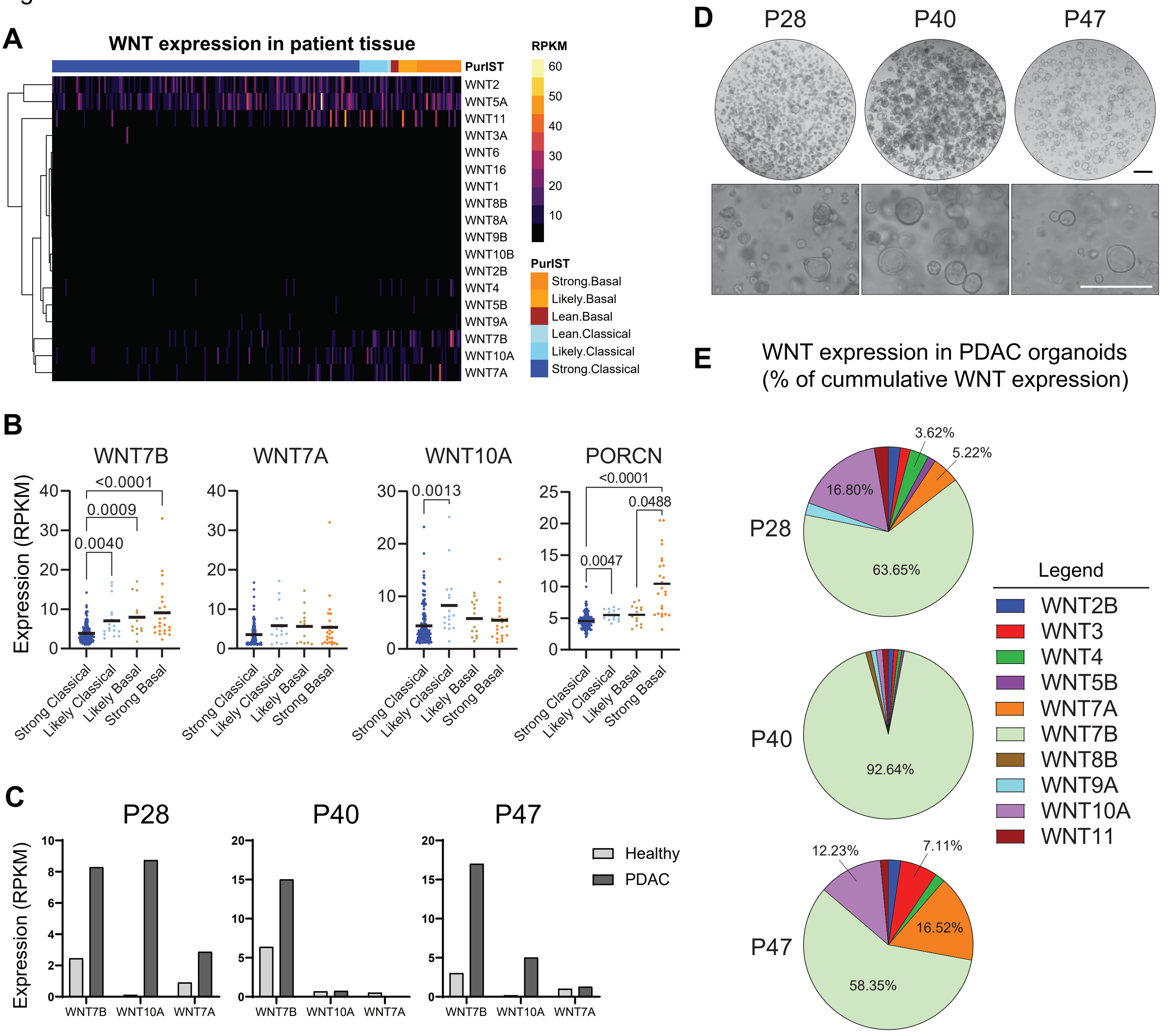
A
WNT expression in patient tissue
PurIST
WNT2
WNT5A
WNT11
WNT3A
WNT6
WNT16
WNT1
WNT8B
WNT8A
WNT9B
WNT10B
WNT2B
WNT4
WNT5B
WNT9A
WNT7B
WNT10A
WNT7A
RPKM
60
50
40
30
20
10
PurIST
Strong.Basal
Likely.Basal
Lean.Basal
Lean.Classical
Likely.Classical
Strong.Classical
B
WNT7B
WNT7A
WNT10A
PORCN
Expression (RPKM)
<0.0001
0.0009
0.0040
0.0013
<0.0001
0.0488
0.0047
Strong Classical
Likely Classical
Likely Basal
Strong Basal
C
P28
P40
P47
Expression (RPKM)
Healthy
PDAC
WNT7B
WNT10A
WNT7A
D
P28
P40
P47
E
WNT expression in PDAC organoids
(% of cummulative WNT expression)
P28
3.62%
5.22%
16.80%
63.65%
P40
92.64%
P47
12.23%
7.11%
16.52%
58.35%
Legend
WNT2B
WNT3
WNT4
WNT5B
WNT7A
WNT7B
WNT8B
WNT9A
WNT10A
WNT11

Figure S2

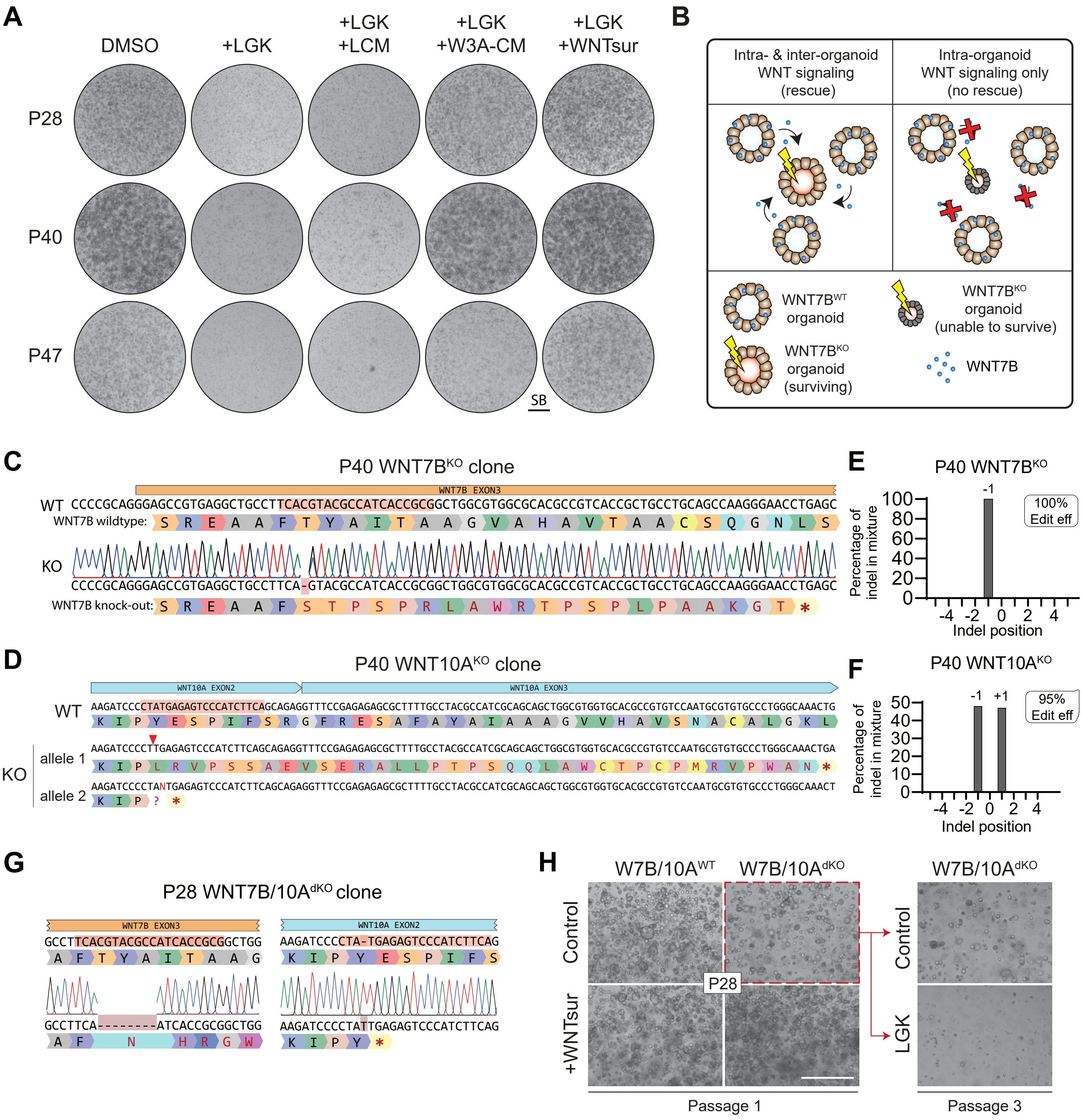
A
DMSO
+LGK
+LGK +LCM
+LGK +W3A-CM
+LGK +WNTsur
P28
P40
P47
SB
B
Intra- & inter-organoid WNT signaling (rescue)
Intra-organoid WNT signaling only (no rescue)
WNT7B^WT organoid
WNT7B^KO organoid (surviving)
WNT7B^KO organoid (unable to survive)
WNT7B
C
P40 WNT7B^KO clone
WNT7B EXON3
WT CCCCGCAGGGAGCCGTGAGGCTGCCTTCACGTACGCCATCACCGCGGCTGGCGTGGCGCACGCCGTCACCGCTGCCTGCAGCCAAGGGAACCTGAGC
WNT7B wildtype:
KO
CCCCGCAGGGAGCCGTGAGGCTGCCTTCA-GTACGCCATCACCGCGGCTGGCGTGGCGCACGCCGTCACCGCTGCCTGCAGCCAAGGGAACCTGAGC
WNT7B knock-out:
E
P40 WNT7B^KO
100% Edit eff
Percentage of indel in mixture
Indel position
D
P40 WNT10A^KO clone
WNT10A EXON2
WNT10A EXON3
WT AAGATCCCCTATGAGAGTCCCATCTTCAGCAGAGGTTTCCGAGAGAGCGCTTTTGCCTACGCCATCGCAGCAGCTGGCGTGGTGCACGCCGTGTCCAATGCGTGTGCCCTGGGCAAACTG
KO
allele 1 AAGATCCCCTTTGAGAGTCCCATCTTCAGCAGAGGTTTCCGAGAGAGCGCTTTTGCCTACGCCATCGCAGCAGCTGGCGTGGTGCACGCCGTGTCCAATGCGTGTGCCCTGGGCAAACTGA
allele 2 AAGATCCCCTANTGAGAGTCCCATCTTCAGCAGAGGTTTCCGAGAGAGCGCTTTTGCCTACGCCATCGCAGCAGCTGGCGTGGTGCACGCCGTGTCCAATGCGTGTGCCCTGGGCAAACT
F
P40 WNT10A^KO
95% Edit eff
Percentage of indel in mixture
Indel position
G
P28 WNT7B/10A^dKO clone
WNT7B EXON3
GCCTTCACGTACGCCATCACCGCGGCTGG
GCCTTCA--------ATCACCGCGGCTGG
WNT10A EXON2
AAGATCCCCTA-TGAGAGTCCCATCTTCAG
AAGATCCCCTATTGAGAGTCCCATCTTCAG
H
W7B/10A^WT
W7B/10A^dKO
W7B/10A^dKO
Control
+WNTsur
P28
Control
LGK
Passage 1
Passage 3

Figure S3

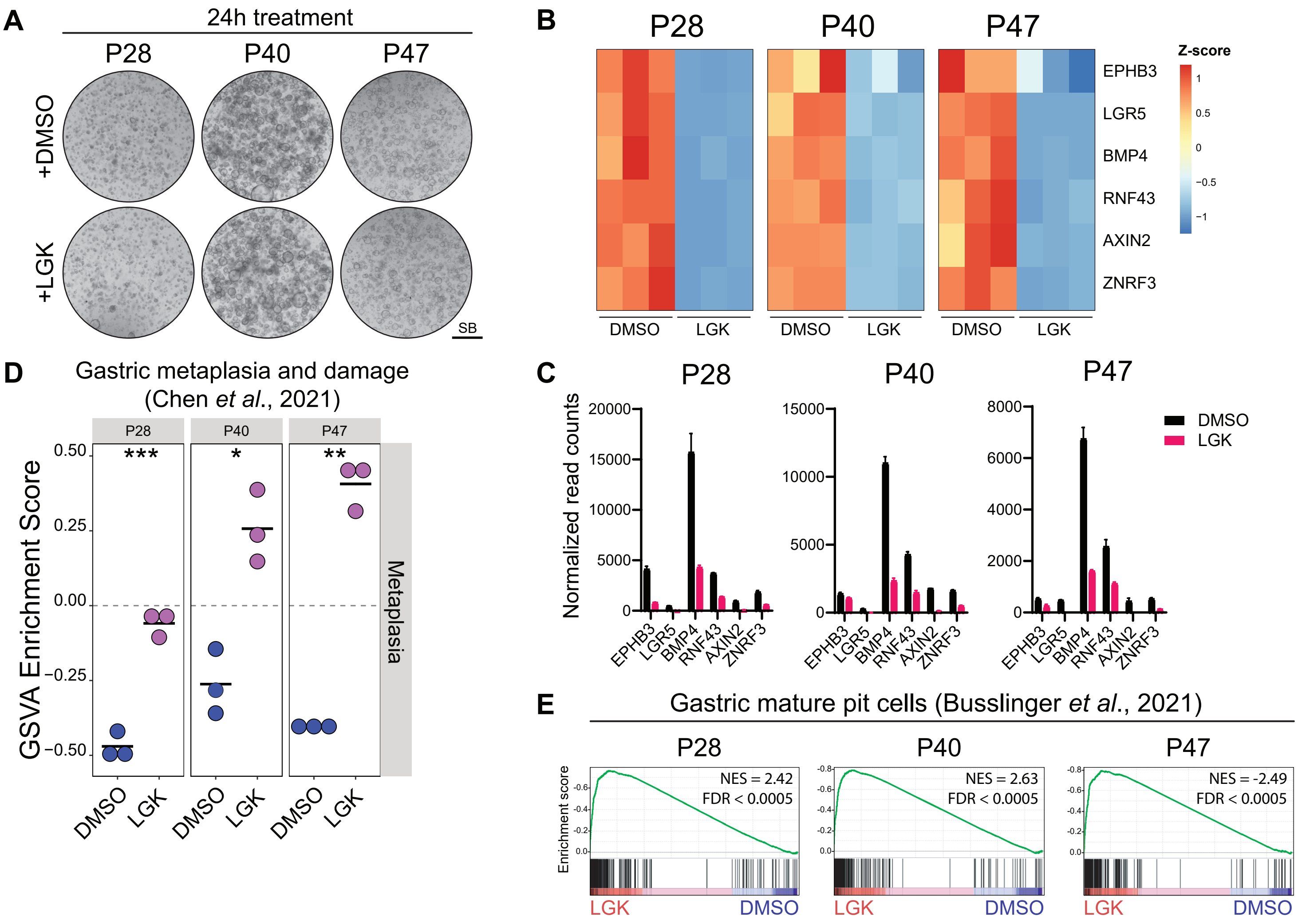
A
24h treatment
P28
P40
P47
+DMSO
+LGK
SB
B
P28
P40
P47
EPHB3
LGR5
BMP4
RNF43
AXIN2
ZNRF3
Z-score
DMSO
LGK
C
P28
P40
P47
Normalized read counts
DMSO
LGK
EPHB3
LGR5
BMP4
RNF43
AXIN2
ZNRF3
D
Gastric metaplasia and damage
(Chen et al., 2021)
GSVA Enrichment Score
***
*
**
Metaplasia
DMSO
LGK
E
Gastric mature pit cells (Busslinger et al., 2021)
Enrichment score
NES = 2.42
FDR < 0.0005
NES = 2.63
FDR < 0.0005
NES = -2.49
FDR < 0.0005
LGK
DMSO

Figure S4

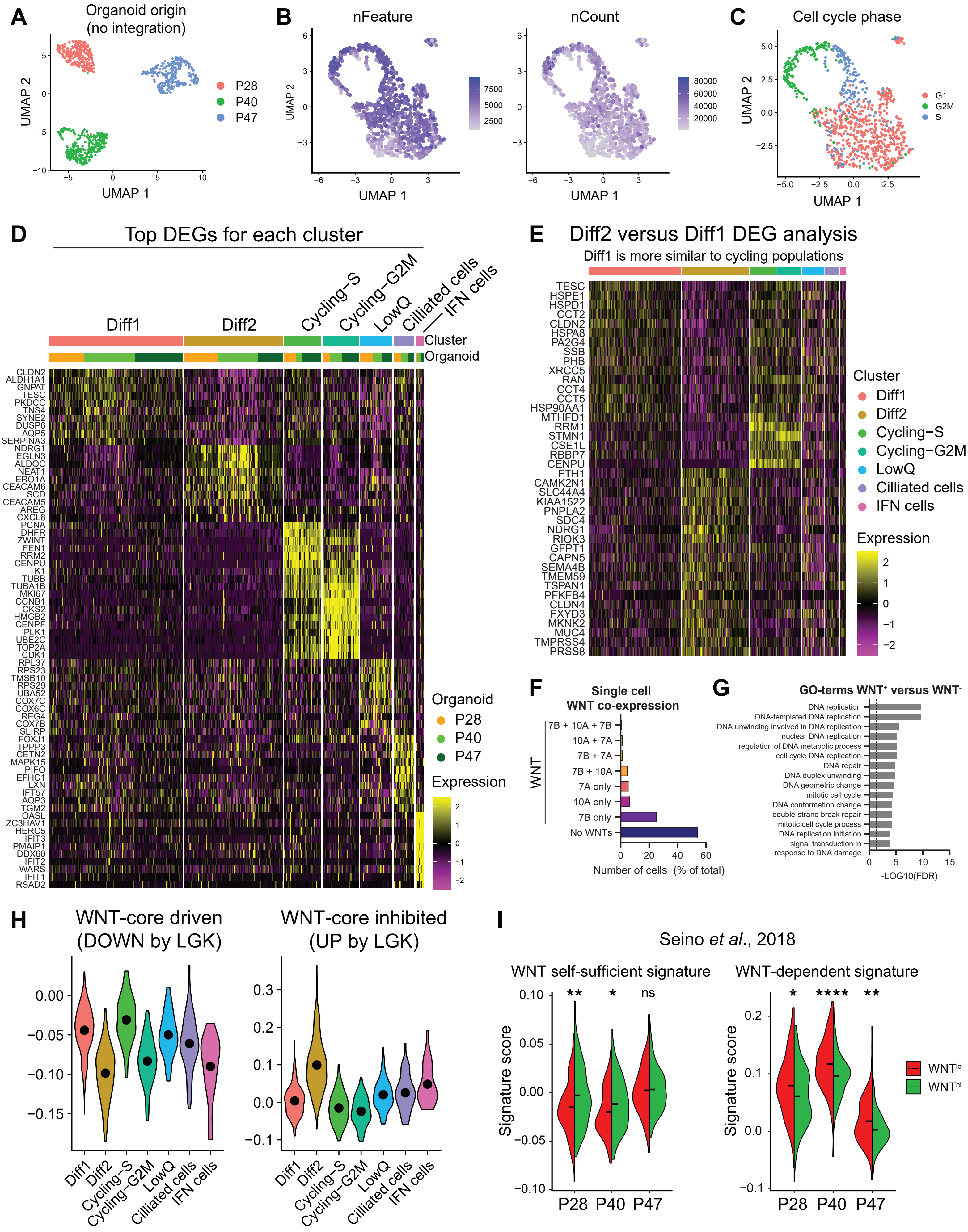
A
Organoid origin
(no integration)
UMAP 2
UMAP 1
P28
P40
P47
B
nFeature
nCount
UMAP 2
UMAP 1
7500
5000
2500
80000
60000
40000
20000
C
Cell cycle phase
UMAP 2
UMAP 1
G1
G2M
S
D
Top DEGs for each cluster
Diff1
Diff2
Cycling−S
Cycling−G2M
LowQ
Ciliated cells
IFN cells
Cluster
Organoid
CLDN2
ALDH1A1
GNPAT
TESC
PKDCC
TNS4
SYNE2
DUSP6
AQP5
SERPINA3
NDRG1
EGLN3
ALDOC
NEAT1
ERO1A
CEACAM6
SCD
CEACAM5
AREG
CXCL8
PCNA
DHFR
ZWINT
FEN1
RRM2
CENPU
TK1
TUBB
TUBA1B
MKI67
CCNB1
CKS2
HMGB2
CENPF
PLK1
UBE2C
TOP2A
CDK1
RPL37
RPS23
TMSB10
RPS29
UBA52
COX7C
COX6C
REG4
COX7B
SLIRP
FOXJ1
TPPP3
CETN2
MAPK15
PIFO
EFHC1
LXN
IFT57
AQP3
TGM2
OASL
ZC3HAV1
HERC5
IFIT3
PMAIP1
DDX60
IFIT2
WARS
IFIT1
RSAD2
Organoid
P28
P40
P47
Expression
E
Diff2 versus Diff1 DEG analysis
Diff1 is more similar to cycling populations
TESC
HSPE1
HSPD1
CCT2
CLDN2
HSPA8
PA2G4
SSB
PHB
XRCC5
RAN
CCT4
CCT5
HSP90AA1
MTHFD1
RRM1
STMN1
CSE1L
RBBP7
CENPU
FTH1
CAMK2N1
SLC44A4
KIAA1522
PNPLA2
SDC4
NDRG1
RIOK3
GFPT1
CAPN5
SEMA4B
TMEM59
TSPAN1
PFKFB4
CLDN4
FXYD3
MKNK2
MUC4
TMPRSS4
PRSS8
Cluster
Diff1
Diff2
Cycling−S
Cycling−G2M
LowQ
Cilliated cells
IFN cells
Expression
F
Single cell
WNT co-expression
WNT
7B + 10A + 7B
10A + 7A
7B + 7A
7B + 10A
7A only
10A only
7B only
No WNTs
Number of cells (% of total)
G
GO-terms WNT+ versus WNT-
DNA replication
DNA-templated DNA replication
DNA unwinding involved in DNA replication
nuclear DNA replication
regulation of DNA metabolic process
cell cycle DNA replication
DNA repair
DNA duplex unwinding
DNA geometric change
mitotic cell cycle
DNA conformation change
double-strand break repair
mitotic cell cycle process
DNA replication initiation
signal transduction in response to DNA damage
-LOG10(FDR)
H
WNT-core driven
(DOWN by LGK)
WNT-core inhibited
(UP by LGK)
Diff1
Diff2
Cycling−S
Cycling−G2M
LowQ
Cilliated cells
IFN cells
I
Seino et al., 2018
WNT self-sufficient signature
WNT-dependent signature
Signature score
**
*
ns
*
****
**
P28
P40
P47
WNTlo
WNThi

Figure S5

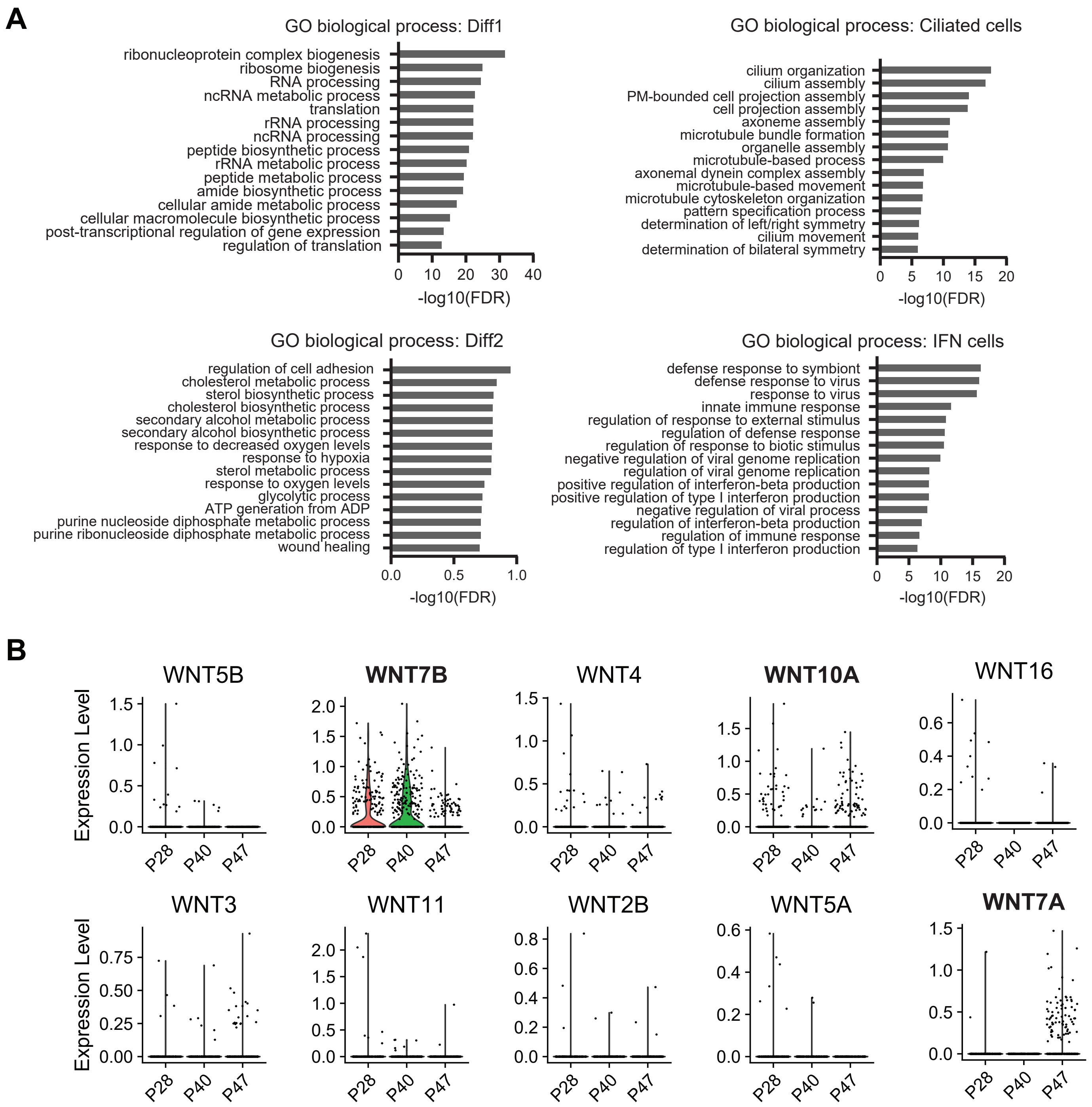

A
GO biological process: Diff1
ribonucleoprotein complex biogenesis
ribosome biogenesis
RNA processing
ncRNA metabolic process
translation
rRNA processing
ncRNA processing
peptide biosynthetic process
rRNA metabolic process
peptide metabolic process
amide biosynthetic process
cellular amide metabolic process
cellular macromolecule biosynthetic process
post-transcriptional regulation of gene expression
regulation of translation
-log10(FDR)
GO biological process: Ciliated cells
cilium organization
cilium assembly
PM-bounded cell projection assembly
cell projection assembly
axoneme assembly
microtubule bundle formation
organelle assembly
microtubule-based process
axonemal dynein complex assembly
microtubule-based movement
microtubule cytoskeleton organization
pattern specification process
determination of left/right symmetry
cilium movement
determination of bilateral symmetry
-log10(FDR)
GO biological process: Diff2
regulation of cell adhesion
cholesterol metabolic process
sterol biosynthetic process
cholesterol biosynthetic process
secondary alcohol metabolic process
secondary alcohol biosynthetic process
response to decreased oxygen levels
response to hypoxia
sterol metabolic process
response to oxygen levels
glycolytic process
ATP generation from ADP
purine nucleoside diphosphate metabolic process
purine ribonucleoside diphosphate metabolic process
wound healing
-log10(FDR)
GO biological process: IFN cells
defense response to symbiont
defense response to virus
response to virus
innate immune response
regulation of response to external stimulus
regulation of defense response
regulation of response to biotic stimulus
negative regulation of viral genome replication
regulation of viral genome replication
positive regulation of interferon-beta production
positive regulation of type I interferon production
negative regulation of viral process
regulation of interferon-beta production
regulation of immune response
regulation of type I interferon production
-log10(FDR)
B
WNT5B
WNT7B
WNT4
WNT10A
WNT16
WNT3
WNT11
WNT2B
WNT5A
WNT7A
Expression Level
P28
P40
P47

Figure S6

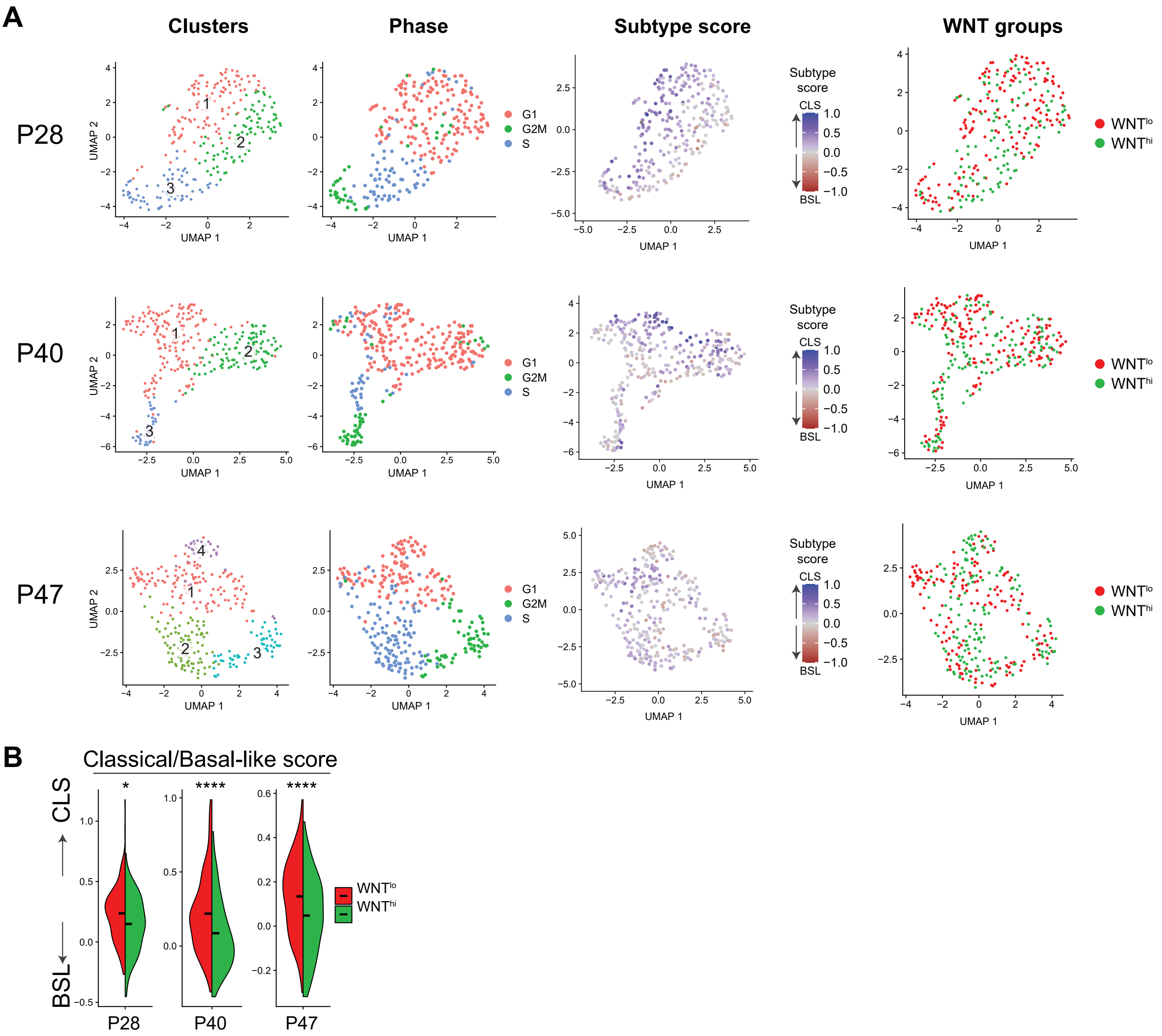

A
Clusters
Phase
Subtype score
WNT groups
P28
P40
P47
UMAP 1
UMAP 2
G1
G2M
S
Subtype score
CLS
BSL
WNTlo
WNThi
B
Classical/Basal-like score
CLS
BSL
*
****
****
P28
P40
P47

Figure S7

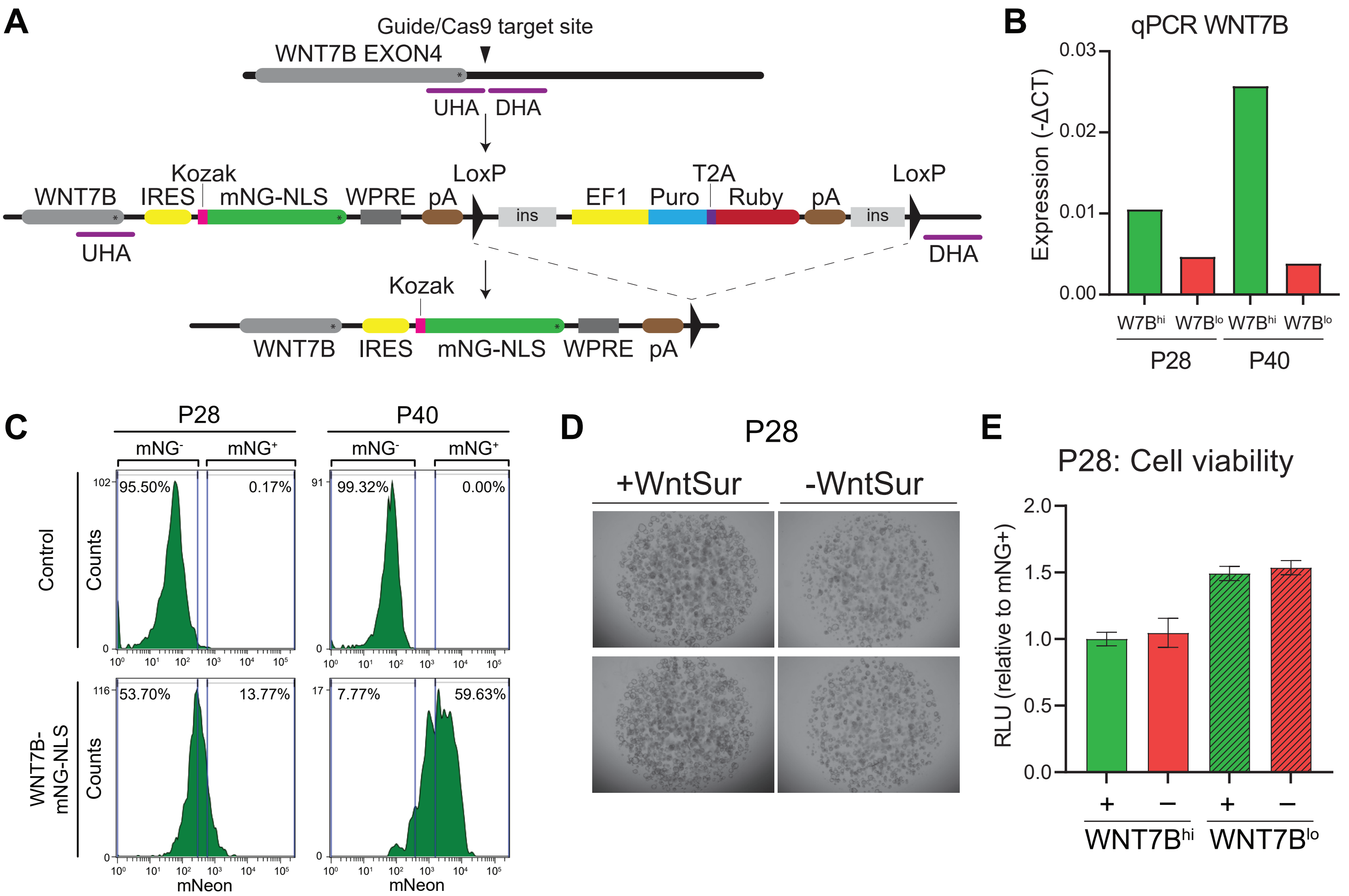
A
Guide/Cas9 target site
WNT7B EXON4
UHA
DHA
Kozak
LoxP
T2A
LoxP
WNT7B
IRES
mNG-NLS
WPRE
pA
ins
EF1
Puro
Ruby
pA
ins
UHA
DHA
Kozak
WNT7B
IRES
mNG-NLS
WPRE
pA
B
qPCR WNT7B
Expression (-ΔCT)
0.03
0.02
0.01
0.00
W7Bhi
W7Blo
W7Bhi
W7Blo
P28
P40
C
P28
P40
mNG-
mNG+
mNG-
mNG+
Control
Counts
95.50%
0.17%
99.32%
0.00%
WNT7B-mNG-NLS
Counts
53.70%
13.77%
7.77%
59.63%
mNeon
mNeon
D
P28
+WntSur
-WntSur
E
P28: Cell viability
RLU (relative to mNG+)
2.0
1.5
1.0
0.5
0.0
+
−
+
−
WNT7Bhi
WNT7Blo

Figure S8

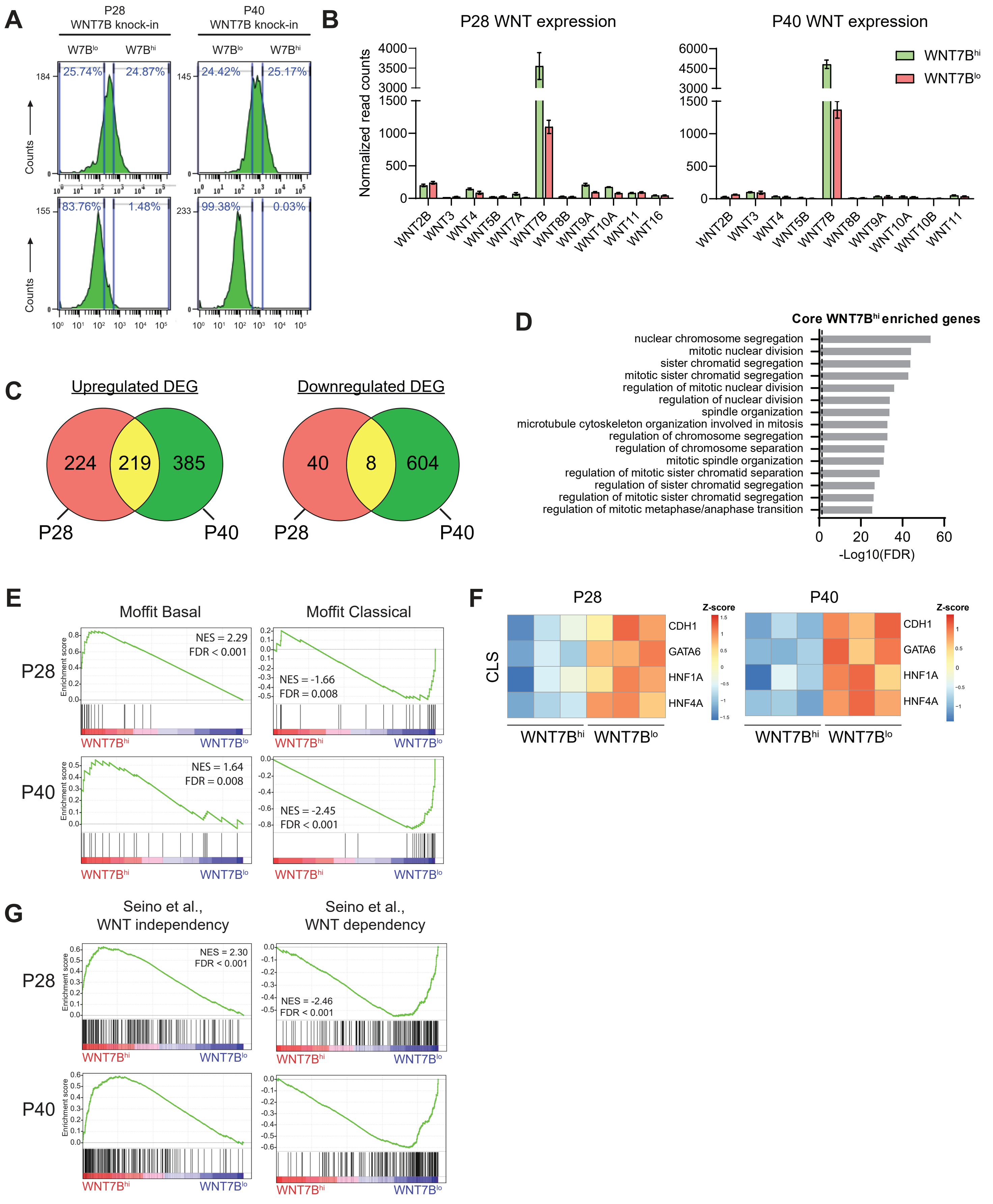
A
P28
WNT7B knock-in
P40
WNT7B knock-in
W7B^lo
W7B^hi
25.74%
24.87%
24.42%
25.17%
83.76%
1.48%
99.38%
0.03%
Counts
B
P28 WNT expression
P40 WNT expression
Normalized read counts
WNT7B^hi
WNT7B^lo
WNT2B
WNT3
WNT4
WNT5B
WNT7A
WNT7B
WNT8B
WNT9A
WNT10A
WNT11
WNT16
WNT10B
C
Upregulated DEG
224
219
385
P28
P40
Downregulated DEG
40
8
604
D
Core WNT7B^hi enriched genes
nuclear chromosome segregation
mitotic nuclear division
sister chromatid segregation
mitotic sister chromatid segregation
regulation of mitotic nuclear division
regulation of nuclear division
spindle organization
microtubule cytoskeleton organization involved in mitosis
regulation of chromosome segregation
regulation of chromosome separation
mitotic spindle organization
regulation of mitotic sister chromatid separation
regulation of sister chromatid segregation
regulation of mitotic sister chromatid segregation
regulation of mitotic metaphase/anaphase transition
-Log10(FDR)
E
Moffit Basal
Moffit Classical
P28
P40
Enrichment score
NES = 2.29
FDR < 0.001
NES = -1.66
FDR = 0.008
NES = 1.64
FDR = 0.008
NES = -2.45
FDR < 0.001
F
P28
P40
CLS
CDH1
GATA6
HNF1A
HNF4A
Z-score
G
Seino et al.,
WNT independency
Seino et al.,
WNT dependency
NES = 2.30
FDR < 0.001
NES = -2.46
FDR < 0.001
